# The Ribosome Maturation Factor Rea1 utilizes nucleotide independent and ATP-hydrolysis driven Linker remodelling for the removal of ribosome assembly factors

**DOI:** 10.1101/2022.11.03.515000

**Authors:** Johan Busselez, Geraldine Koenig, Torben Klos, Piotr Sosnowski, Nils Marechal, Hugo Gizardin-Fredon, Sarah Cianferani, Carine Dominique, Yves Henry, Anthony Henras, Helgo Schmidt

## Abstract

The ribosome maturation factor Rea1 (or Midasin) catalyses the removal of assembly factors from large ribosomal subunit precursors to promote their export from the nucleus to the cytosol. Rea1 consists of nearly 5000 amino-acid residues and belongs to the AAA+ protein family. It consists of a ring of six AAA+ domains from which the ≈ 1700 amino-acid residue linker emerges that is subdivided into stem, middle and top domains. A flexible and unstructured D/E rich region connects the linker top to a MIDAS (metal ion dependent adhesion site) domain, which is able to bind the assembly factor substrates. Despite its key importance for ribosome maturation, the Rea1 mechanism driving assembly factor removal is still poorly understood. Here we demonstrate that the Rea1 linker is essential for assembly factor removal. It rotates and swings towards the AAA+ ring following a complex remodelling scheme involving nucleotide independent as well as nucleotide dependent steps. ATP-hydrolysis is required to engage the linker with the AAA+ ring and ultimately with the AAA+ ring docked MIDAS domain. The interaction between the linker top and the MIDAS domain allows force transmission for assembly factor removal.

## Introduction

Ribosomes synthesise proteins and the production of ribosomes is the most energy consuming process in cells. In eukaryotes more than 200 assembly factors are involved in the production of ribosomes [1]. Ribosome assembly starts in the nucleolus with the transcription of the 5S and 35S rRNAs by RNA polymerases III and I, respectively. The 35S rRNA is subsequently cleaved into a smaller 20S and a larger 27S part. The 20S rRNA forms the basis of the future 40S ribosomal subunit, while the 27S rRNA associates with various ribosomal proteins, ribosomal assembly and maturation factors as well as the 5S rRNA to form pre-60S ribosomal (pre60S) particles [2]. These particles are subsequently exported to the cytosol via the nucleoplasm. During this process, they are gradually transformed into mature 60S subunits by transiently interacting with additional assembly factors that promote conformational rearrangements and rRNA processing steps [2]. The 60S assembly intermediates have been extensively studied using *Saccharomyces cerevisiae* as a model organism [3, 4]. One of the earliest assembly intermediates identified in the nucleolus is a pre60S particle that carries the Ytm1 complex, consisting of Ymt1, Erb1 and Nop7 [5] and involved in the processing of the 27S rRNA [6]. To promote transfer of this particle to the nucleoplasm, the Ytm1 complex has to be removed, a reaction that is carried out by Rea1 (also known as “Ylr106p” or “Midasin”) [7, 8]. Rea1 associates with pre60S particles via its interaction with the Rix1 complex, which consists of Rix1, lpi1 and lpi3 [9, 10]. The Rea1 catalysed removal of the Ytm1 complex is essential for the nucleolar export of pre60S particles, because disrupting the interaction between Rea1 and Ytm1 leads to the accumulation of pre60S particles in the nucleolus [8].

In the nucleoplasm, pre60S particles bind to the assembly factor Rsa4, which tightly associates with another assembly factor, Nsa2. Nsa2 wraps around the H89 rRNA helix of the immature peptidyltransferase centre [11–13]. In addition to the Ytm1 complex, Rea1 also mechanically removes Rsa4 from pre60S particles [10]. In doing so, it might indirectly force Nsa2 to pull on the H89 rRNA helix to drag this important architectural feature of the peptidyltransferase centre into its correct position [11].

The Rea1 mediated Rsa4 removal is also crucial for the export of pre60S particles from the nucleoplasm to the cytosol. The removal of Rsa4 leads to an activation of the GTPase activity of Nug2 [14]. Subsequently, Nug2-GDP dissociates, which allows the nuclear export factor Nmd3 to bind to the former Nug2 site. Nmd3 in turn associates with Crm1 and RanGTP to trigger the export of pre60S particles to the cytosol [14]. Disrupting the Rea1-Rsa4 interaction leads to the accumulation of pre60S particles in the nucleoplasm and severe growth defects in *S. cerevisiae* [10], demonstrating the importance of Rea1 also for nucleoplasmic pre60S particle export.

Rea1 is conserved from yeast to mammals and related to the motor protein dynein [15]. Its deletion in yeast leads to non-viable strains [15, 16] underscoring the essential role of this complex molecular machine, which consists of nearly 5000 amino-acid residues. Rea1 folds into a hexameric ring of AAA+ (<u>A</u>TPases <u>a</u>ssociated with various cellular <u>a</u>ctivities) domains, each divided into large (AAAL) and small sub domains (AAAS). From the hexameric AAA+ ring a ≈ 1700 amino-acid linker domain emerges, consisting of stem-, middle- and top subdomains [17] (Figure 1A and B). The linker top2 domain connects to a ≈ 600 amino-acid glutamate and aspartate rich region which ends in a ≈ 300 amino-acid metal ion dependent adhesion site (MIDAS) domain [15, 17–19] (Figure 1A and B). The MIDAS domain interacts with the ubiquitin like (UBL) domains of Ytm1 and Rsa4 [8, 10]. The interaction is mediated by a Mg^2+^ ion and resembles the integrin MIDAS – ligand interaction [10, 20]. The MIDAS domain is able to dock onto the AAA+ ring [17, 19] and in the context of the Rea1-pre60S particle complex this docking site places the MIDAS domain in direct contact with the Rsa4 UBL domain [18]. Recently, the catch bond character of the MIDAS-UBL domain interaction has been demonstrated [21].

**Figure 1:**
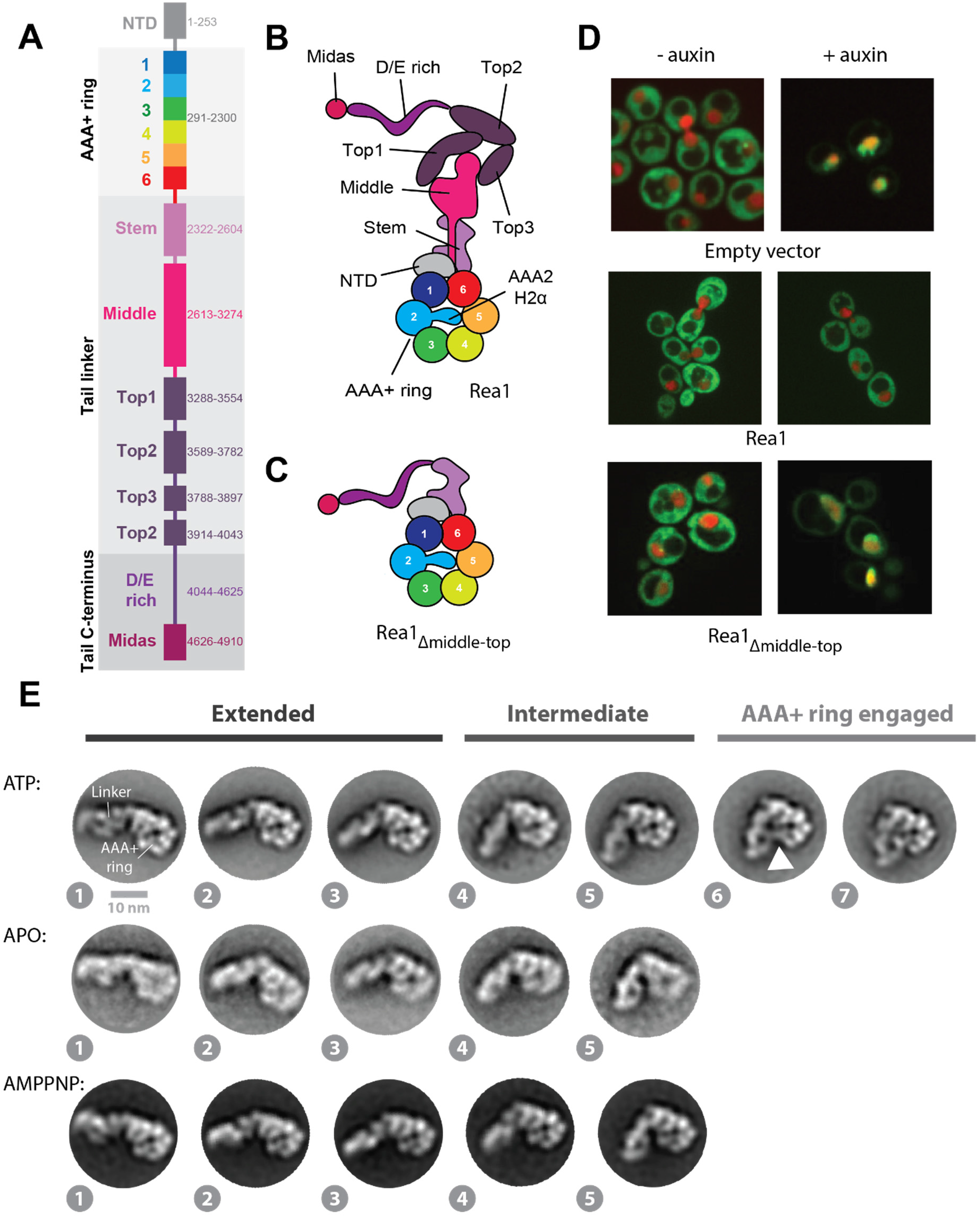
The Rea1 linker is a functionally important structural element and shows nucleotide independent as well as nucleotide dependent remodelling. **A.** Domain organization of Rea1. **B.** Schematic cartoon representation of Rea1. The helix 2 α-helical insertion of the AAA2 domain (AAA2H2α) sits in the central pore of the AAA+ ring. **C.** Schematic cartoon representation of a construct lacking the linker middle and top domains. The flexible D/E rich region with the substrate binding MIDAS domain are directly fused to the linker stem domain. **D.** Yeast nuclear export assay of pre-ribosomal particles. The pre-ribosomal particle marker Rpl25 is fused to GFP. Histone-mCherry marks the nucleus. The endogenous Rea1 is under the control of the auxin degron system. Upper panels: The addition of auxin leads to the accumulation of GFP fluorescence in the nucleus indicating a pre60S nuclear export defect due to degraded endogenous Rea1. Middle panels: The export defect can be rescued by a plasmid harbouring a Rea1_wt_ copy. Lower panels: Providing a plasmid harbouring the construct in C. does not rescue the export defect, suggesting the linker middle and top domains are functionally important. **E.** Negative stain 2D classes representing AAA+ ring top views of Rea1_wt_ in the presence of ATP, absence of nucleotide as well as in the presence of AMPPNP. States 1 – 5 represent the extended and intermediate linker conformations, which do not require nucleotide. In contrast to states 1 – 5, the AAA+ ring engaged states 6 and 7 require ATP hydrolysis. The white arrow head highlights a connection between the linker top and the AAA+ ring.

The question how force is transmitted to the substrate engaged MIDAS domain to remove Rsa4 or Ytm1 from pre60S particles has been controversial. Ytm1 and Rsa4 *in-vitro* release assays have established that the force production for assembly factor removal from pre60S particles requires ATP-hydrolysis [8, 10], but the Rea1 conformations associated with ATP-hydrolysis have not been described yet. Early hypotheses suggested the Rea1 linker and MIDAS domain would move as a rigid body during ATP-hydrolysis to remove assembly factors from pre60S particles [10]. However, recent high-resolution cryoEM studies did not detect nucleotide dependent Rea1 linker remodelling [17, 19]. It has been suggested that linker remodelling might not be relevant for assembly factor removal. Instead is was proposed that nucleotide driven conformational rearrangements in the AAA+ ring are directly communicated to the AAA+ ring docked, substrate engaged MIDAS domain to produce the force for assembly factor removal [19, 22].

Here we provide evidence for the functional importance of the Rea1 linker region and demonstrate nucleotide independent as well as ATP-hydrolysis dependent linker remodelling in Rea1. We show that linker remodelling consists of two main components, a rotation of the linker middle and top domains relative to the linker stem domain and a pivot movement towards the AAA+ ring. Furthermore, we demonstrate that linker remodelling is able to produce mechanical force. In the final remodelling step, the linker top interacts with the AAA+ ring docked MIDAS domain allowing direct transmission of force for assembly factor removal. Our results reveal key mechanistic events of one of the most complex eukaryotic ribosome maturation factors, whose mode of action has remained elusive since the first description of Rea1 20 years ago.

## Results

### The Rea1 linker top and middle domains are functionally important

To investigate the functional relevance of the Rea1 linker region, we created a Rea1 construct with deleted linker top and middle domains and fused the D/E rich region with the attached MIDAS domain directly to the linker stem (Rea1Δ_middle-top_) (Figure 1C). We tested the functionality of Rea1Δ_middle-top_ using a modified version of a *Saccharomyces cerevisiae* GFP-Rpl25 pre60S export assay [10]. In this assay, the ability of a Rea1 construct to promote the nuclear export of pre60S particles is monitored by the occurrence of GFP fluorescence in the cytoplasm. We tagged the endogenous Rea1 gene with the auxin degron system, provided a plasmid encoding Rea1Δ_middle-top_ and monitored the cellular GFP fluorescence distribution after auxin induced endogenous Rea1 degradation. Consistent with a crucial functional role of the linker top and middle domains in pre60S particle nuclear export, we observe a lack of GFP fluorescence in the cytoplasm and GFP-fluorescence accumulation in the nucleus (Figure 1D).

### The Rea1 linker undergoes nucleotide independent as well as nucleotide dependent remodelling

Having established the functional importance of the linker region, we decided to investigate the nucleotide depended conformational changes of the Rea1 linker. Recently, we [17] and other groups [19] aimed at the determination of 3D structures of distinct linker conformations by electron microscopy (EM), but were not able to visualize linker remodelling. We reasoned that such an approach might fail to detect alternative linker conformations if they are in low abundance and/or classify into a limited number of 2D views thereby precluding the calculation of interpretable 3D EM maps.

We decided to analyse Rea1 linker remodelling by negative stain electron microscopy (EM). Compared to cryoEM, negative stain EM offers the advantage of increased signal-to-noise ratios for individual particles, which should ease the detection of low abundance Rea1 linker conformations. We reasoned that even though the resolution of negative stain EM is limited, large-scale remodelling events of the ≈200 kDa linker domain should still be detectable. We also limited our image processing workflow at the 2D classification stage to avoid failing to detect alternative linker conformation during 3D classification due to insufficient 2D projection distributions.

First, we characterized wild-type *S. cerevisiae* Rea1 (Rea1_wt_) in the presence of ATP. We were able to obtain several 2D classes with identical top view onto the AAA+ ring. In these 2D classes, the linker samples a range of different conformations with respect to the AAA+ ring as illustrated in Figure 1E. We tentatively sorted them into “Extended”, “Intermediate” and “AAA+ ring engaged” linker conformations” and assigned the numbers 1 - 7 from the most extended to the most compact Rea1 linker state (Figure 1E). States 6 and 7, which represent the AAA+ ring engaged linker conformations are clearly minority classes representing only ≈ 0.8% of the total particles in the data set. States 1 – 3 of the extended and state 4 of the intermediate linker conformations are similar to Rea1 linker conformations described in earlier studies [10] (Supplementary figure 1). In the nucleotide free APO data set, the AAA+ ring top view particles exclusively sorted into the extended and intermediate linker conformations (states 1 - 5) (Figure 1E), suggesting that the AAA+ ring engaged linker conformations require the presence of nucleotide. To further investigate whether ATP binding or hydrolysis is needed to engage the linker with the AAA+ ring, we collected a data set in the presence of the non-hydrolysable ATP analogue AMPPNP. Again, only the extended and intermediate linker conformations were sampled (states 1 - 5) (Figure 1E) indicating that linker AAA+ ring engagement requires ATP-hydrolysis.

Next, we investigated a Rea1 mutant lacking the AAA2 helix 2 α-helical insert (AAA2H2α) (Rea1_ΔAAA2H2α_) (compare Figure 1B). In a previous study we demonstrated that AAA2H2α is an auto-inhibitory regulator of the Rea1 ATPase activity that also prevents the docking of the MIDAS domain onto the AAA+ ring [17]. The relocation of AAA2H2α from the central pore of the AAA+ ring and the docking of the MIDAS domain onto the AAA+ ring is also observed when Rea1_wt_ is bound to Rsa4-pre60S particles [18, 23]. This suggests that the Rea1_ΔAAA2H2α_ mutant resembles Rea1_wt_ when bound to pre60S particles.

We determined negative stain EM 2D class averages of the ATP, APO, ADP and AMPPNP states. In the presence of ATP, the AAA+ ring top view particles sorted again in extended, intermediate as well as AAA+ ring engaged classes (Figure 2). The linker conformations are highly similar to the ones observed for Rea1_wt_ (Figure 1E), suggesting that linker remodelling is not altered in Rea1_ΔAAA2H2α_. In the absence of nucleotide, we detect linker conformations that are consistent with states 1 – 5 of the extended and intermediate classes (Supplementary figure 2). However, the AAA+ ring was disordered suggesting that in Rea1_ΔAAA2H2α_ nucleotide is required to stabilize the AAA+ ring. Incubating Rea1_ΔAAA2H2α_ with ADP or AMPPNP restricts linker remodelling to the extended and intermediate classes (Figure 2) indicating that - like in the case of Rea1_wt_-ATP-hydrolysis is required for linker AAA+ ring engagement. AAA+ ring engaged classes are again minority views representing only ≈0.4% of the total particles.

**Figure 2:**
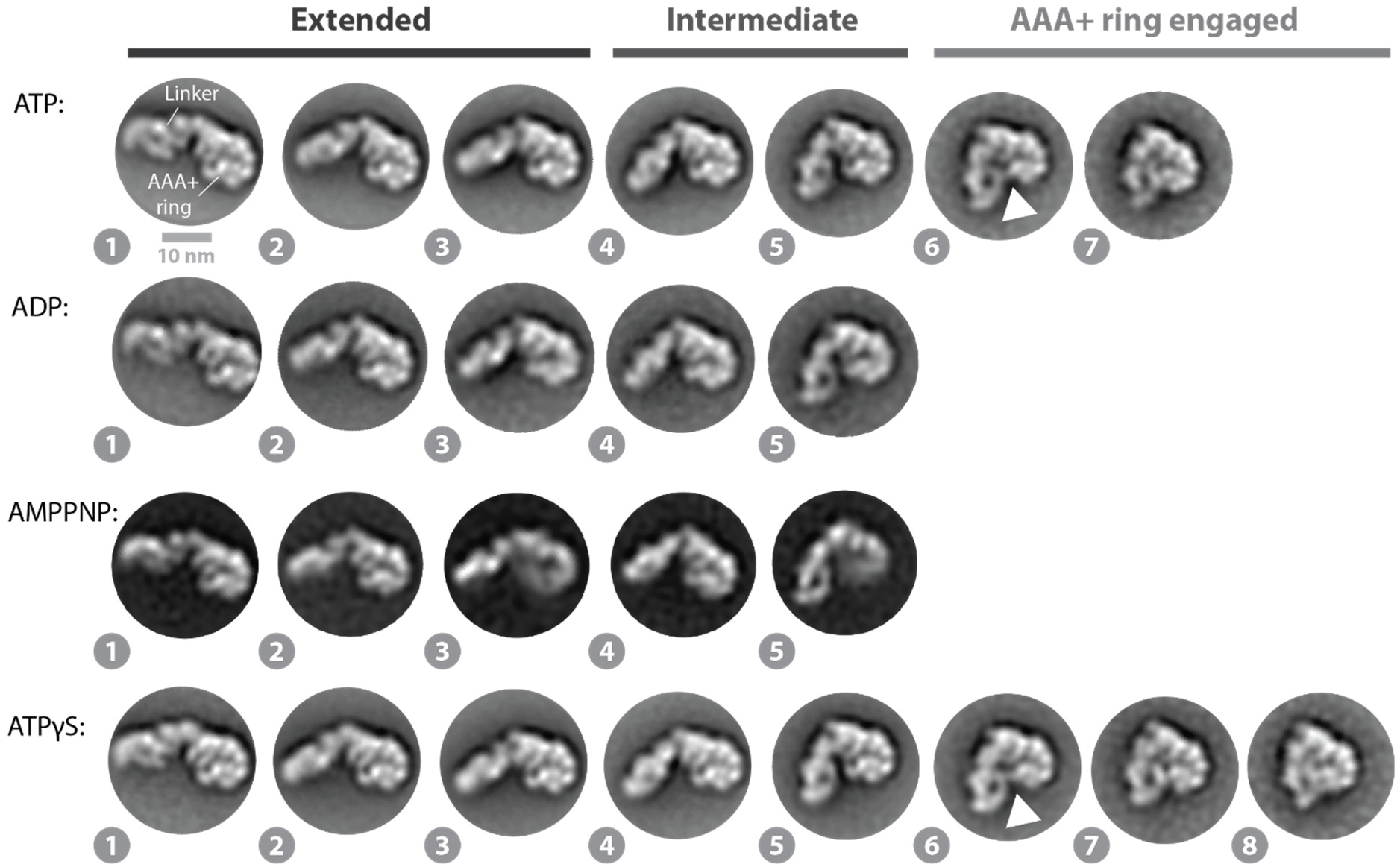
Linker remodelling in the Rea1_ΔAAA2H2α_. The analysis of the ATP, ADP, AMPPNP and ATPγS data sets indicates that linker remodelling in Rea1_ΔAAA2H2α_ is highly similar to linker remodelling in Rea1_wt_. Like in the case of Rea1_wt_, ATP-hydrolysis is required to engage the linker with the AAA+ ring. Unlike Rea1_wt_, Rea1_ΔAAA2H2α_ is able to sample state 8 in the presence of the slowly hydrolysable ATP analogue ATPγS. White arrow heads highlight a connection between the linker top and the AAA+ ring.

Recent cryoEM studies on a AAA+ unfoldase made use of the slowly hydrolysable ATP analogue ATPγS to enrich transient protein conformations [24]. We tested if ATPγS might stabilize the AAA+ ring engaged linker states in Rea1_ΔAAA2H2α_. Although we did not observe a substantial enrichment (≈ 0.4% ATP vs. ≈ 1.5% ATPγS), we detected state 8, an additional AAA+ ring engaged linker conformation (Figure 2). We also investigated, if state 8 is sampled in Rea1_wt_ in the presence of ATPγS. We detected 2D class averages similar to state 8 of Rea1_ΔAAA2H2α_ but with less well-defined structural features of the linker indicating increased structural flexibility (Supplementary figure 3).

Collectively, these results demonstrate that distinct Rea1 linker remodelling steps exist. The extended and intermediate linker conformations (states 1 – 5) are being sampled even in the absence of any nucleotide highlighting the intrinsic conformational flexibility of the linker. The AAA+ ring engagement of the linker requires ATP-hydrolysis, which suggests an important functional role for these low abundance conformations. This view is also supported by Rsa4 and Ymt1 *in-vitro* release assays, which demonstrated that only ATP but not AMPPNP enables Rea1 to catalyse the removal of assembly factor substrates from pre60S particles [8, 10]. These findings indicate that the extended and intermediate linker conformations (states 1 – 5) (Figures 1E and 2) are insufficient to support functionality and that additional conformations linked to ATP-hydrolysis (states 6-8) have to be sampled to catalyse assembly factor removal. Rea1_ΔAAA2H2α_ largely resembles linker remodelling in Rea1_wt_, but - unlike Rea1_wt_ - is able to stably sample state 8.

### The Linker top and middle domains rotate and pivot towards the AAA+ ring docked MIDAS domain during remodelling

Since Rea1_ͤAAA2H2α_ in the ATPγS state revealed the highest number of linker remodelling conformations, we thought to annotate the Rea1 subdomains in the corresponding 2D class averages to analyse Rea1 linker remodelling in more detail. To this end, we determined a Rea1_ΔAAA2H2α_ ATPγS 3D cryoEM structure (Figure 3A, B and supplementary figure 4) to generate 2D projections for the subdomain assignment in our negative stain EM 2D classes. Consistent with previous cryoEM investigations [17], we could only obtain an interpretable 3D reconstruction showing the linker in the straight conformation, which was the dominant structural state in the sample and already described in earlier work (Supplementary figure 5) [17–19].

**Figure 3:**
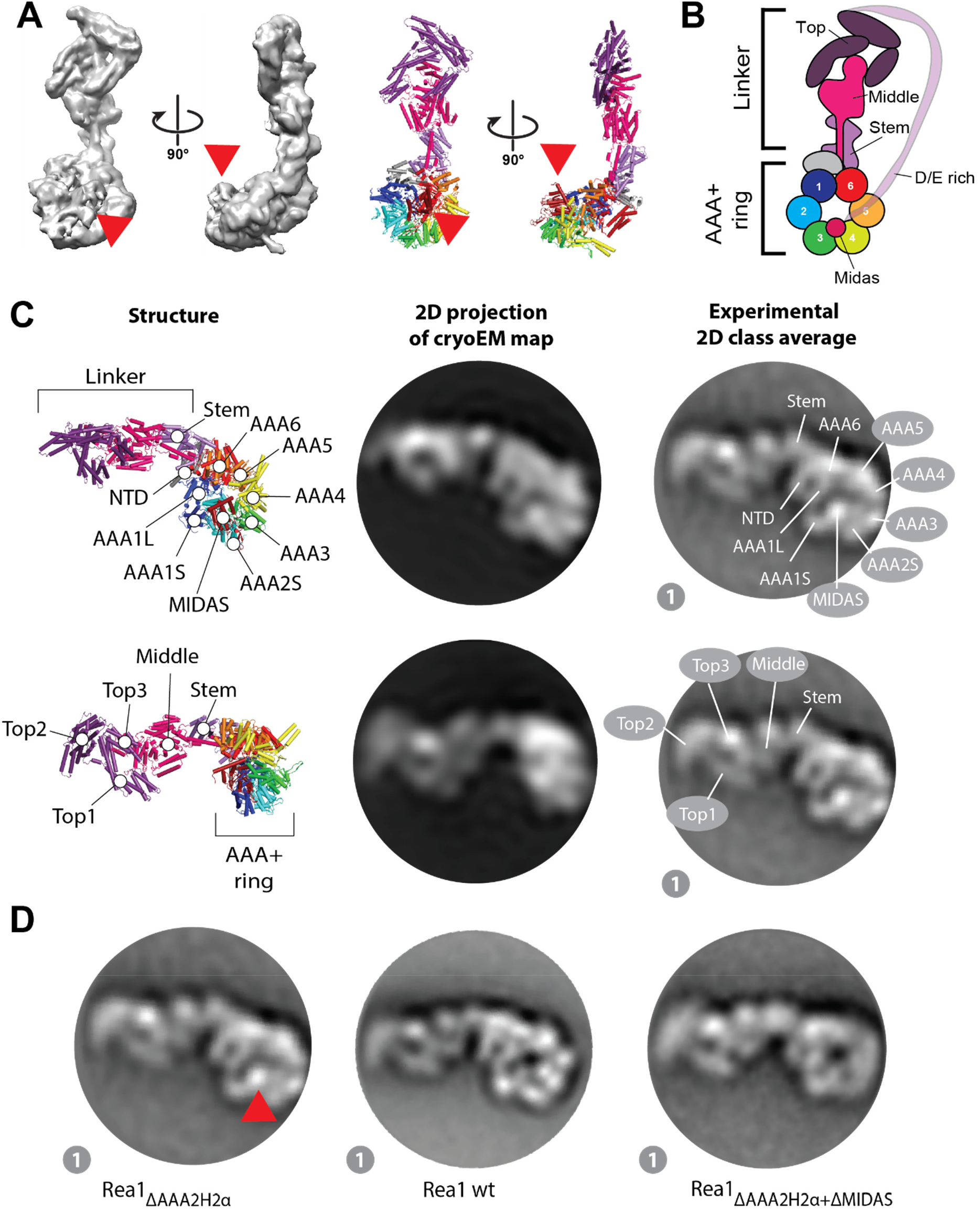
Domain assignments in negative stain 2D class averages of Rea1_ΔAAA2H2α_. **A.** CryoEM map (left panels) and cartoon representation (right panels) of Rea1_ΔAAA2H2α_ in the presence of ATPγS. The red arrow heads highlight the AAA+ ring docked MIDAS domain. **B.** Schematic cartoon representation of the structure in A. The α-helical extension of AAA2 normally occupying the centre of the pore (compare Figure 1B) has been deleted, which allows the MIDAS domain to dock onto the AAA+ ring. The D/E rich region connecting the MIDAS domain to the linker top is flexible and not visible in the structure. **C.** Upper panels: A 2D projection of the cryoEM map in A. low pass filtered to 25 Å shows a good match for the AAA+ ring region in the state 1 negative stain 2D class average. The projection allows the assignment of the NTD, AAA1 – AAA6, the linker stem and the MIDAS domain. In contrast to the AAA+ ring region, the linker top and middle domains adopt a different conformation from the one seen in state 1. Lower panels: With a different 2D projection of the low pass filtered cryoEM map a good match for the linker region in the state 1 negative stain 2D class average can be produced allowing the assignment of the linker top1, top2 and top3 domains as well as the linker middle domain. The AAA+ ring does not match up with the AAA+ ring in the state 1 negative stain 2D class average. This mismatch indicates that – compared to the cryoEM structure in A. - the linker top and middle domains in state 1 have moved with respect to the AAA+ ring. **D.** The assignment of the MIDAS domain in state 1 of Rea1_ΔAAA2H2α_ (red arrow head, left panel) is further supported by comparisons with state 1 of Rea1_wt_, where the MIDAS domain is absent from the AAA+ ring (middle panel, compare also supplementary figure 7) as well as analysis of the Rea1_ΔAAA2H2α+ΔMIDAS_ double mutant (right panel).

We generated a 2D projection from the cryoEM map with high similarity to the AAA+ ring in our negative stain EM 2D class averages (Figure 3C), but the linker part did not resemble any of the observed linker conformations of the eight states (Figure 2). Using a different orientation, we were able to obtain a 2D projection matching the linker of state 1 but not the AAA+ ring (Figure 3C). The combination of both projections allowed us to assign the NTD, the six AAA+ domains of the AAA+ ring as well as the linker stem, middle and top domains and the MIDAS domain in state 1 (Figure 3C). Our analysis also demonstrates that the linker in state 1 has already undergone a rigid-body movement with respect to the AAA+ ring compared to the straight linker conformation in our cryoEM structure. We approximate that the linker top and middle domains have rotated ≈ 30 ° counter clock wise around the long linker axis and bent ≈ 45 ° towards the plane of the AAA+ ring (Supplementary figure 6).

A prominent feature of the Rea1_ΔAAA2H2α_ AAA+ ring in states 1 – 8 is a bright spot on the AAA+ ring, which we interpret as AAA+ ring docked MIDAS domain (Figure 3C). This feature is absent in the Rea1_wt_ classes (Figure 3D and supplementary figure 7), which is consistent with earlier findings demonstrating that the presence of the AAA2 H2 α-helical insert in Rea1_wt_ interferes with the AAA+ ring docking of the MIDAS domain [17]. We also directly confirmed our MIDAS domain assignment by analysing the Rea1_ΔAAA2H2α-ΔMIDAS_ double mutant (Figure 3D).

Next, we aligned the eight states onto their AAA+ rings (Figure 4A, Movie S1). These AAA+ ring alignments suggest that the linker top and middle domains pivot towards the AAA+ ring docked MIDAS domain during linker remodelling. The pivot point during this movement is located between the linker middle and stem domains. These alignments further suggest that the linker top and middle domains rotate during the pivot movement. To better visualize this additional transformation, we aligned states 1 - 8 on the long linker axis (Figure 4B, Movie S2) to demonstrate the rotation, which occurs during states 1 - 5. The linker top and middle domains behave approximately as a rigid body during the rotation as suggested by a series of 2D projections of the linker top and middle domains rotated around the long linker axis (Figures 4C and D). We estimate the rotation angle between states 1 and 5 to be ≈ 100 ° (Figure 4E). In the final linker remodelling conformation, state 8, the linker top2 and top3 domains are in close proximity to the AAA+ ring docked MIDAS domain (Figure 4A and B, Movies S1 and S2).

**Figure 4:**
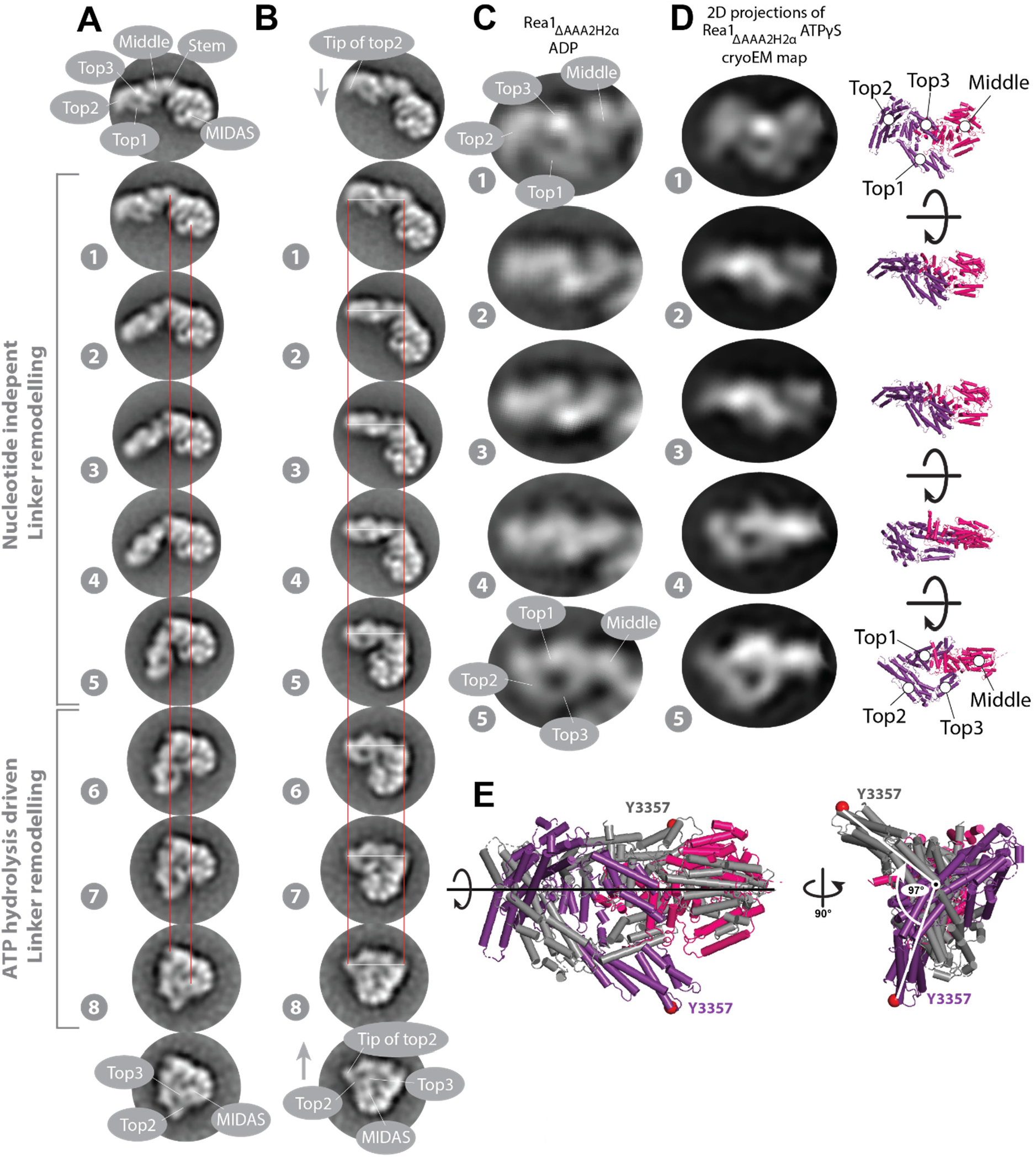
The Rea1 linker pivots and rotates during remodelling. **A.** States 1 – 8 of Rea1_ΔAAA2H2α_ ATPγS aligned on the linker stem and MIDAS domains (red lines). The linker middle and top domains swing towards the AAA+ docked MIDAS domain (compare also movie S1). The region between the linker stem and middle domains acts as pivot point. **B.** States 1 – 8 of Rea1_ΔAAA2H2α_ ATPγS aligned on long linker axis (tip of linker top2 domain and linker stem domain, white and red lines). The linker top and middle domains rotate during the pivot swing (compare also movie S2). The tip of the linker top2 domain points downwards in state 1 (grey arrow) but upwards in state 8 (grey arrow) highlighting the rotation. In the final linker remodelling conformation, state 8, the linker top2 and top3 domains as well as the MIDAS domain are in close proximity. States 1 - 5 were also observed under APO conditions (compare Figure 1E) indicating that large parts of the linker swing and the linker rotation are nucleotide independent and are part of the intrinsic conformational flexibility of the linker. The engagement of the rotated linker with the AAA+ ring during states 6 – 8 requires ATP hydrolysis. **C.** Enlarged views of linker region in states 1 – 5 of Rea1_ΔAAA2H2α_ ADP. **D.** Left panels: Series of 2D projections of the linker top-middle part of the Rea1_ΔAAA2H2α_ ATPγS cryoEM map low pass filtered to 25 Å and rotated around the long linker axis. Right panels: corresponding structures. The rotation of the linker in A. and B. can be approximated by a rigid-body rotation of the linker top and middle domains around the long linker axis. Additional internal rearrangements of the linker middle and top domains with respect to each other cannot be excluded. **E.** Aligning states 1 (color coded) and 5 (grey) of D. on the long linker axis indicates a total rotation angle of ≈ 100 °. Equivalent Cα atoms are shown as red spheres.

These results reveal that the linker top and middle domains undergo a complex series of movements with respect to the linker stem domain and the AAA+ ring during linker remodelling. Compared to the straight linker conformation in Figure 3A, they rotate and pivot towards the plane of the AAA+ ring in an initial movement to reach state 1. From the position in state 1 they pivot towards the AAA+ ring docked MIDAS domain and further rotate around the long linker axis. The rotation largely happens during states 1-5, which cover the extended and intermediate linker conformations. Since these states are also observed under APO conditions (Figure 1E and supplementary figure 2), we conclude that the rotational movement and the additional swing towards the AAA+ ring do not require energy suggesting they are part of the intrinsic conformational flexibility of the Rea1 linker. The energy of ATP-hydrolysis is needed to engage the fully rotated linker top and middle domains with the AAA+ ring during states 6 - 8 of the AAA+ ring engaged linker conformations. The connection between the linker and the AAA+ ring in state 6 that separates the AAA+ ring engaged conformations from the intermediate conformations occurs between the linker top3 domain and AAA1S (Supplementary figure 8). In state 8, the linker top2 and top3 domains are in close proximity to the AAA+ ring docked MIDAS domain (Figure 4A and B). The fact that state 8 was not observed in Rea1_wt_, which does not show the AAA+ ring docked MIDAS domain (Figure 3D and supplementary figure 7), suggests that the presence of the MIDAS domain at the AAA+ ring is essential for sampling state 8.

### The Linker top interacts with the MIDAS domain and linker remodelling is a force producing event

Since the linker top2 and top3 domains in state 8 are located next to the MIDAS domain, we wanted to test if there is a direct interaction. To this end, we carried out crosslinking mass spectrometry on the Rea1_ΔAAA2H2α_ mutant in the presence of ATPγS (Supplementary figure 9). Although our negative stain EM analysis suggests that state 8 only represents ≈ 0.2% of the total particles in our sample, we were able to detect two crosslinks between K3955 in the top2 domain and either K4662 or K4668 in the MIDAS domain (Figure 5A). An additional crosslink was detected between K3569, located in the linker top1-top2 loop region that associates with the top2 domain [17], and K4662. The K3569-K4662, K3955-K4662 and K3955-K4668 crosslinks further support our domain assignments in state 8 (Figure 4A and B) and hint at a direct interaction between the linker top and the MIDAS domain. Consistent with our negative stain EM analysis, we did not detect these crosslinks in the presence of AMPPNP.

**Figure 5:**
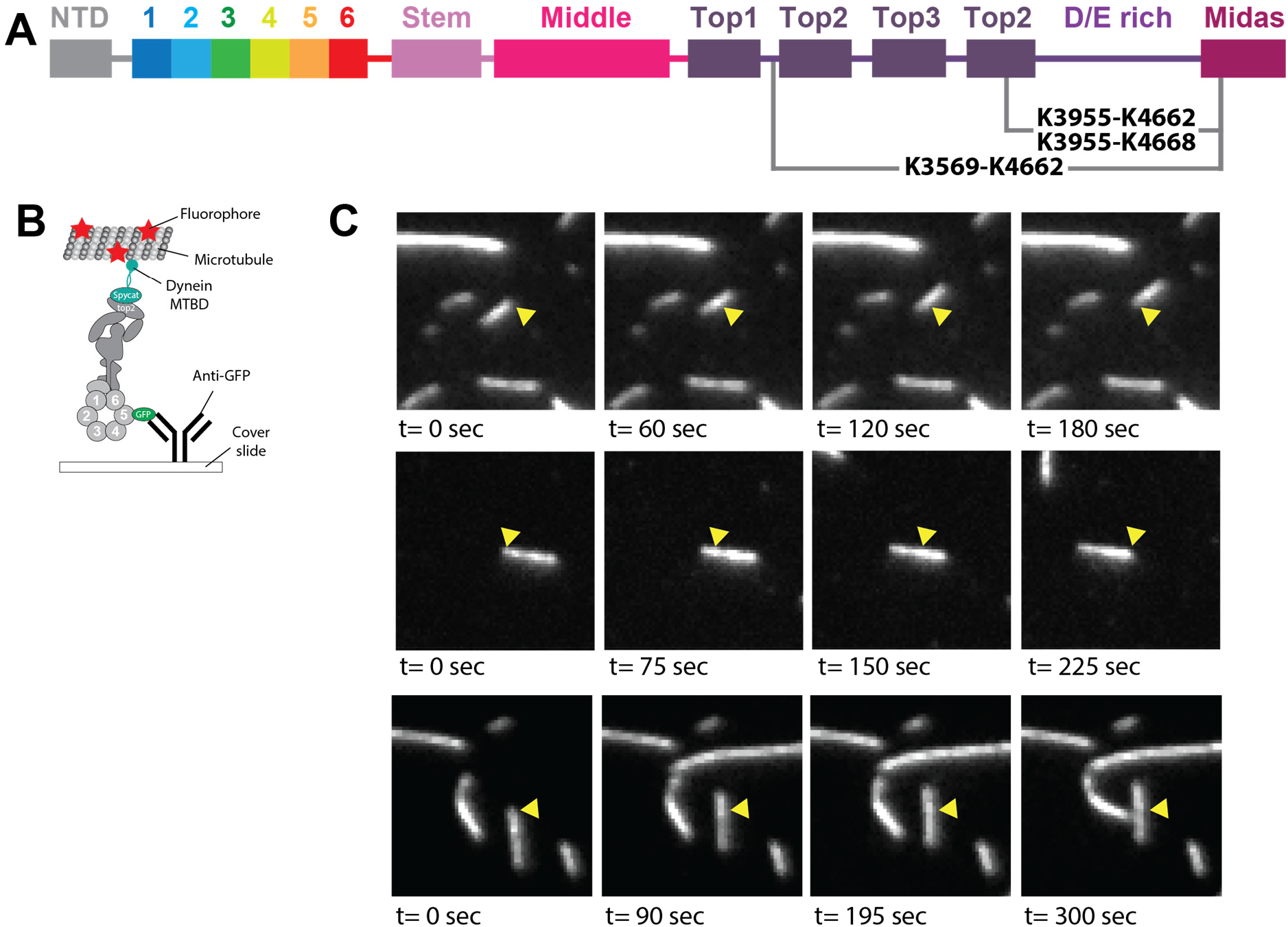
The linker top is able to interact with the MIDAS domain and linker remodelling is a force producing event. **A.**Three crosslinks supporting state 8 detected in Rea1_ΔAAA2H2α_ in the presence of ATPγS. The K3569-K4662, K3955-K4662 and K3955-K4668 crosslinks suggest a direct interaction between the linker top2 domain and a highly conserved loop region in the MIDAS domain (also compare Figure 4A and B). **B**. We carried out microtubule gliding assays with a Rea1_ΔAAA2H2α_ construct to provide evidence that Rea1 linker remodelling produces force. The dynein microtubule binding domain (cyan) was fused to the linker top2 domain using the spycatcher/spytag approach and GFP to AAA5. The construct was anchored to a cover slide via GFP-antibodies and fluorescently labelled microtubules we applied. **C**. Four movie frames of three microtubule gliding events. The yellow arrow heads mark the position of the microtubule at the beginning of the movie (also compare movies S3-S5). The microtubule gliding events suggest that the remodelling of the linker with respect to the AAA+ ring is able to produce mechanical force.

The top2 domain crosslink partners K4662 and K4668 are located in the highly conserved E4656-K4700 loop of the MIDAS domain. This loop region harbours an NLS sequence that is required for the nuclear import of Rea1 [20]. It was also demonstrated that the E4656-K4700 loop is essential for assembly factor removal. Deleting and replacing it with an alternative NLS sequence rescued nuclear import but prevented Rea1 from removing its Rsa4 assembly factor substrate from pre60S particles [20].

To provide additional support for an interaction between the linker top and the MIDAS domain, we carried out yeast-two-hybrid assays, but the expression of the fusion constructs turned out to be toxic for the cells (Supplementary figure 10A). However, a subsequent immunoprecipitation of the MIDAS domain construct provided evidence for an interaction with the linker top2/top3 domains (Supplementary figure 10B).

In order to probe if linker remodelling is able to produce mechanical force, we carried out total-internal-reflection-fluorescence (TIRF) microscopy based microtubule gliding assays. We worked with the Rea1_ΔAAA2H2α_ background since this construct shows elevated ATPase activity compared to the wt [17]. We anticipated that the flexible, unstructured and highly negatively charged D/E rich region (theoretical pI: 3.77) of Rea1 might prevent the binding of microtubules and truncated ≈ 80% of the D/E-rich region as well as the whole MIDAS domain. We further modified Rea1_ΔAAA2H2α+Δ4168-4907_ by fusing a GFP to AAA5 and a spytag into the linker top2 domain. The GFP allowed us to anchor the construct to anti-GFP decorated cover slides and we covalently fused a spycatcher construct carrying the dynein microtubule binding domain into the linker top2 domain via the spytag-spycatcher interaction [25] (Figure 5B). After the application of fluorescently labelled microtubules, we screened for microtubule gliding events. Since force production for microtubule gliding will rely on energy provided by ATP-hydrolysis and our negative stain EM analysis suggests that only a minority of Rea1 molecules adopt ATP-hydrolysis associated conformations in the presence of ATP, microtubule gliding is expected to be a rare event. Consistent with this assumption, we observe occasional events of slow, directed microtubule gliding over several minutes (Figure 5C, Movies S3-S5) indicating that the remodelling of the linker top with respect to the AAA+ ring is able to produce mechanical force.

Taken together, these results suggest that the mechanical force produced by linker remodelling might be directly applied to the AAA+ring-docked MIDAS domain via the linker top2/top3 domains to remove assembly factors from pre60S particles.

### The Linker middle domain acts as a crucial hub for linker remodelling

Next, we wanted to establish which structural elements of the linker are crucial for its remodelling. In a previous study, we identified an α-helical extension of the linker middle domain that contacts the Rea1 AAA+ ring [17] (Figure 6A). We speculated that this α-helical extension might participate in the communication of ATP induced conformational changes from the AAA+ ring into the linker to drive its remodelling [17]. In order to probe its functional role, we deleted this α-helix (Rea1Δ2916-2974) and characterized linker remodelling in the presence of ATP by negative stain EM. Our analysis reveals that Rea1Δ2916-2974 is able to sample states 1-6 (Figure 6B). There is no evidence for state 7, which we would expect to detect under these nucleotide conditions (compare Figure 1E), suggesting that this mutant is impaired in its ability to sample the linker states normally associated with ATP-hydrolysis.

**Figure 6:**
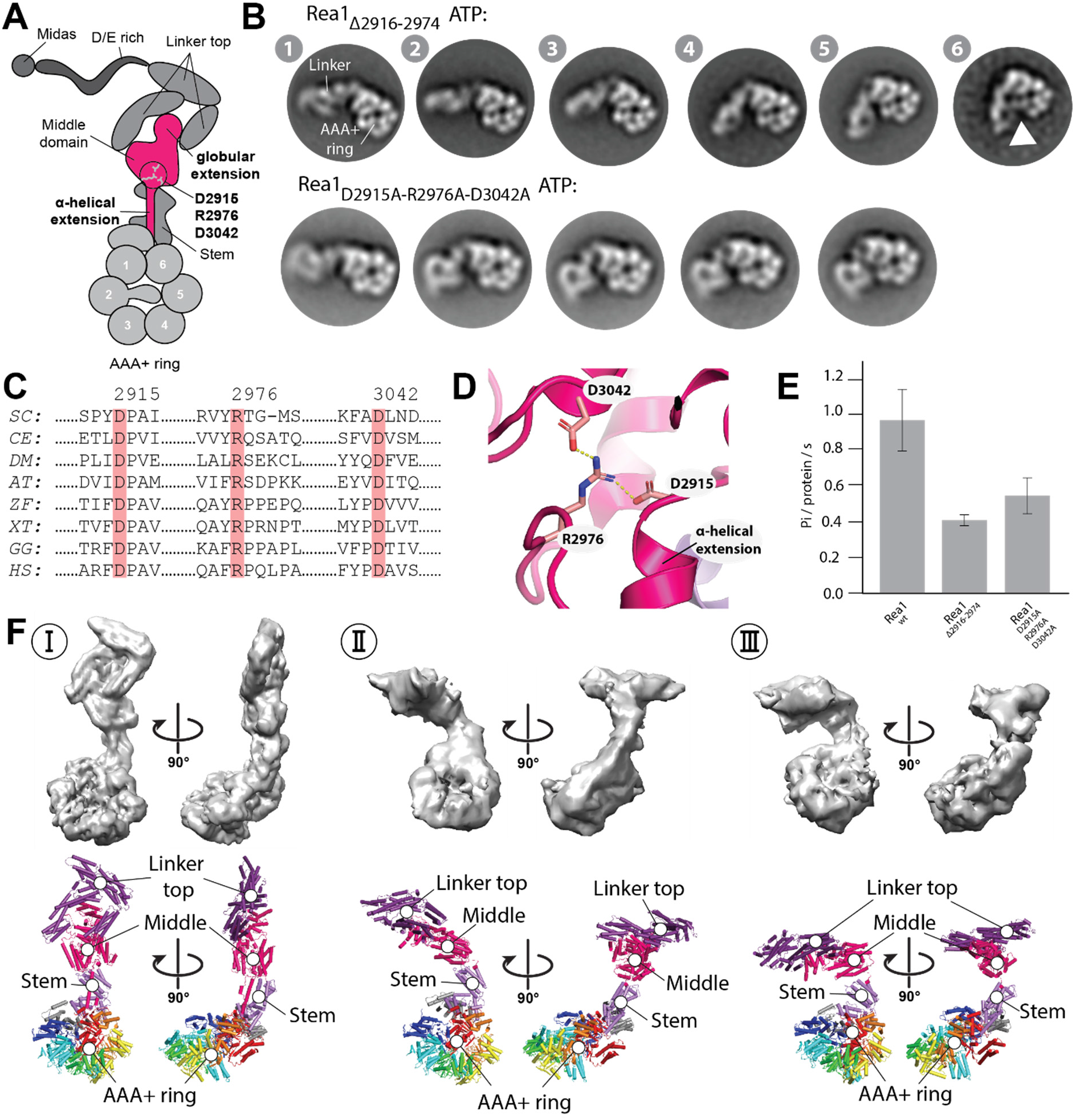
The linker middle domain has essential functions during linker remodelling. **A.** Schematic representation of three structural elements in the linker middle domain. The α-helical extension (aa 2916-2974), the globular extension (aa 3072-3244) and the highly conserved salt-bridge network D2915-R2976-D3042. **B.** Deleting the α-helical extension or disrupting the D2915-R2976-D3042 saltbridge network results in altered linker remodelling pathways. Deleting the α-helical extension prevents the sampling of state 7. The disruption of the D2915-R2976-D3042 still allows the linker swing towards the AAA+ ring but prevents the rotation of the linker top and middle domains around the long linker axis as well as the subsequent engagement with the AAA+ ring (compare also movie S6). Deleting the globular extension leads to the truncation of the whole linker (see supplementary figure 11) indicating that it serves a critical role for the stability of the linker. **C.** The D2915-R2976-D3042 saltbridge network is conserved in Rea1/Midasin of *SC*: Saccharomyces cerevisiae, *CE*: Caenorhabditis elegans, *DM*: Drosophila melanogaster, *AT*: Arabidopsis thaliana, *ZF*: Zebra fish, *XT*: Xenopus tropicalis *GG*: Gallus gallus, *HS*: Homo sapiens. The *S. cerevisiae* Rea1 amino-acid residue numbering is shown at the top of the multiple sequence alignment. **D.** The D2915-R2976-D3042 salt-bridge network in the near-atomic resolution model of the *S. cerevisiae* Rea1 linker (PDB-ID: 6hyd; Sosnowski et al., 2018). The dashed yellow lines indicate hydrogen bonds with a distance of 2.7 Å. **E.** The inability of the Rea1 linker top and middle domains to correctly engage with the Rea1 AAA+ ring affects the ATPase activity. Deleting the α-helical extension (aa 2916-2974) or disrupting the highly conserved D2915-R2976-D3042 salt-bridge network in the linker middle domain reduce the Rea1 ATPase activity by ≈ 50%. The compromised ATPase activity indicates that the AAA+ ring engagement of the linker top and middle stimulates the Rea1 ATPase activity. **F.** Three cryoEM structures of the Rea1D2915A-R2976A-D3042A mutant in the presence of ATP. The linker conformation I is highly similar to the linker conformation of Rea1_ΔAAA2H2α_ ATPγS (compare Figure 3A and supplementary figure 5E). In Conformations II and III the linker top and middle domains have swung towards the AAA+ ring (compare also supplementary figure 13). The region between the linker middle and stem domain acts as pivot point.

Another prominent feature of the Rea1 linker middle domain is the globular extension that supports the U-shaped arrangement of the linker top [17] (Figure 6A). Deleting the globular extension in the construct Rea1_Δ3072-3244_ destabilizes the linker top and leads to partial Rea1 degradation so that only the Rea1 AAA+ ring can be detected in our negative stain analysis (Supplementary figure 11).

Next, we screened for highly conserved amino-acid residues in the Rea1 linker. Although the general conservation within the Rea1 linker domain is low, we nevertheless identified a highly conserved salt-bridge network, D2915-R2976-D3042, in close proximity to the α-helical extension of the linker middle domain (Figure 6A, C and D). The disruption of this salt-bridge network by alanine mutations changed the remodelling pathway of the linker. While the region between the linker middle and stem domains still acts as pivot point for the swing of linker top and middle domains towards the AAA+ ring, Rea1_D2915A-R2976A-D3042A_ did not show the rotation around the long linker axis that we consistently detected in all previous data sets (Figure 6B, movie S6). Consequently, the linker is not able to engage with the AAA+ ring. The inability of Rea1_Δ2916-2974_ and Rea1_D2915A-R2976A-D3042A_ linker to correctly engage with the AAA+ ring in the presence of ATP also affects the ATPase activity of these constructs (Figure 6E). These defects suggest that linker engagement with the AAA+ ring in Rea1_wt_ stimulates the ATPase activity.

We also characterized Rea1_D2915A-R2976A-D3042A_ by cryoEM in the presence of ATP. We were able to obtain three reconstructions at medium to low resolution (Figure 6F and supplementary figure 12). As expected, the straight linker, was the dominant class (conformation I) (Supplementary figure 5E). It refined to a resolution sufficient to resolve secondary structure elements. In addition, we were able to resolve two alternative linker conformations of Rea1_D2915A-R2976A-D3042A_ (conformations II. and III.) at a resolution allowing the docking of the linker stem-AAA+ ring and the linker top-middle domains. As expected, conformations II. and III. were related by a swing of the linker top and middle domains towards the AAA+ ring without rotation around the long linker axis (Supplementary figure 13). These 3D cryoEM reconstructions confirm our initial assignment of the linker middle-stem region as pivot point for linker remodelling.

To assess the functionality of the Rea1_Δ2916-2974_ and Rea1_D2915A-R2976AD3042A_ mutants, we first tested their ability to support growth in the absence of endogenous wild type Rea1. Centromeric plasmids directing expression from the *REA1* promoter of these Rea1 mutants were transformed in a heterozygous *REA1/rea1::kanR* diploid strain bearing a deletion of one *REA1* alelle. Following sporulation and tetrad dissection, haploid spores expressing Rea1_Δ2916-2974_ were unable to form colonies (Figure 7A), indicating that the α-helical extension of the linker middle domain is essential for viability. In contrast, *rea1::kanR* haploids expressing Rea1_D2915A-R2976AD3042A_ could be obtained. However, they are very slow growing, underscoring the functional importance of the conserved salt-bridge network. We next assessed the effects of these deletions and mutations on pre-rRNA processing by northern analyses (Figure 7B). In the case of the lethal Rea1_Δ2916-2974_ construct, this analysis was performed using a *GAL::rea1* strain conditionally expressing endogenous Rea1, transformed with the plasmid mentioned above. The strain was first propagated in galactose-containing medium and shifted to glucose-containing medium to repress wild-type Rea1 expression. The northern analyses indicate that the Rea1_Δ2916-2974_ and Rea1_D2915A-R2976AD3042A_ mutants lead to similar pre-rRNA processing defects as the lack of Rea1, characterized by an increase in the levels of early 35S pre-rRNA and intermediate 27SB pre-rRNA, reflecting defective assembly of both early and intermediate pre-ribosomal particles (Figure 7B and supplementary figure 14). For Rea1_D2915A-R2976AD3042A_, an additional, more thorough analysis was carried out with the haploid *rea1::kanR* strain expressing the mutant. The salt bridge mutations induce an increase in the levels of the 7S pre-rRNA, the precursor to 5.8S rRNA, suggesting that later pre-60S particle maturation stages are also affected (Figure 7C). As a result of the pre-rRNA processing defects, 25S rRNA levels are down in Rea1_D2915A-R2976AD3042A_ (Figure 7C).

**Figure 7:**
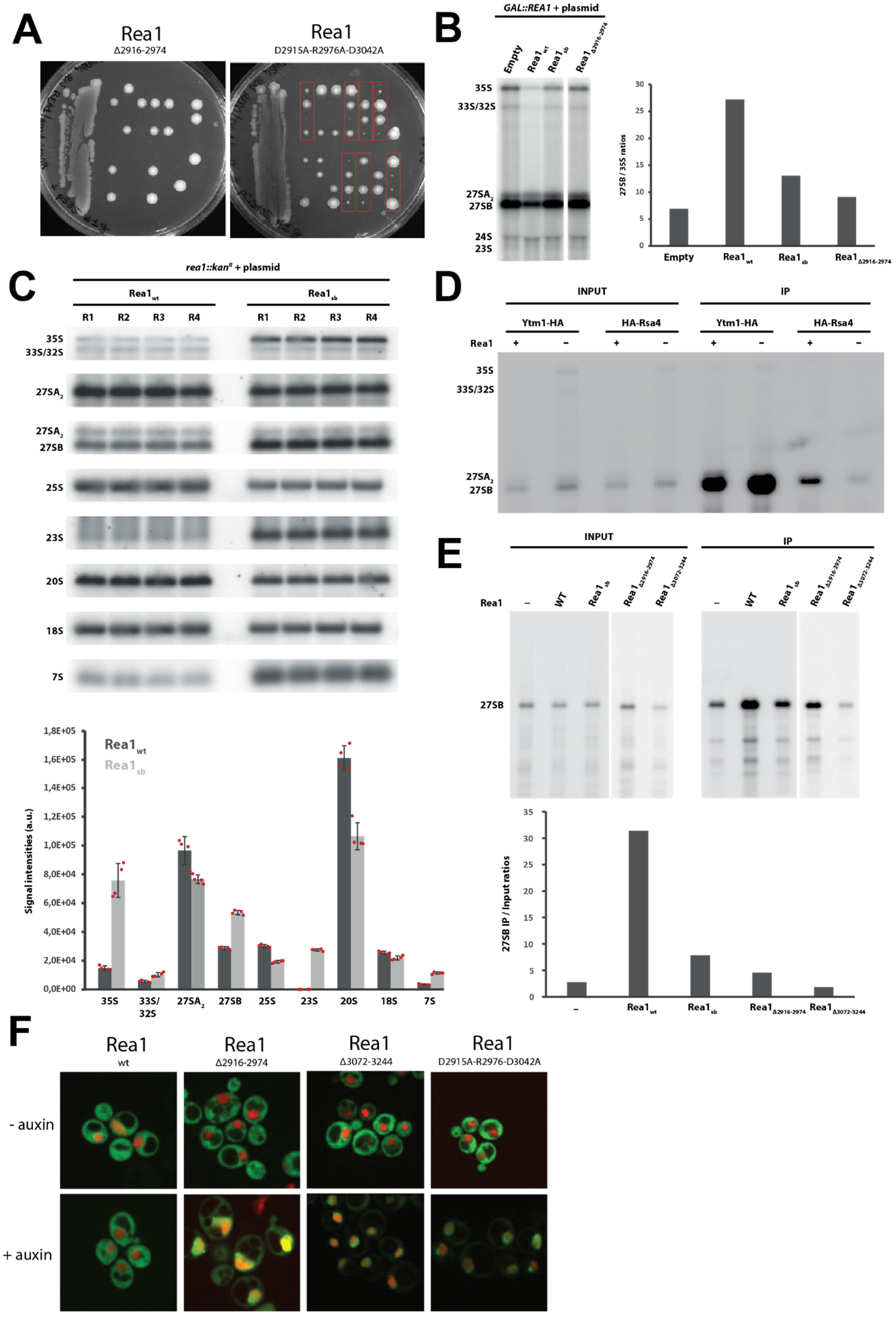
Functional impact of Rea1 linker middle domain deletions and mutations. **A.** Impact of REA1 mutations on yeast viability. A heterozygous REA1/rea1::kanR diploid strain was transformed with plasmids directing expression of the indicated Rea1 mutants. After sporulation of the resulting strains, tetrads were dissected and haploid spores were spotted in rows on YPD medium. Only the spores containing the Rea1_D2915A-R2976A-D3042A_ mutant (Rea1sb) could support viability (red rectangles), although growth was strongly impaired. **B.** Impact of Rea1 mutations on pre-rRNA processing. Left panel: Northern analyses of pre-rRNA processing in GAL::rea1 cells expressing the indicated Rea1 mutants (“Rea1_sb_” stands for Rea1_D2915A-R2976A-D3042A_) or wild-type Rea1 (Rea1_wt_) from plasmids or transformed with an empty vector (EV), shifted 24 hours on glucose-containing medium. Right panel: Quantification of 27SB/35S ratio. The mutants display reduced ratios compared to the Rea1_wt_ control indicative of pre60S maturation defects. **C.** Upper panel: Detailed Northern pre-rRNA processing analysis of rea1::kanR strain expressing Rea1_D2915A-R2976A-D3042A_ (“Rea1_sb_”) from a plasmid. Four wt strains and four Rea1_D2915A-R2976A-D3042A_ expressing strains were analysed in parallel. Detection of the indicated (pre-)rRNAs by northern hybridization with anti-sense oligonucleotide probes. Lower panel: levels of the indicated (pre-)rRNAs, or pre-rRNA ratios in the indicated mutants or wild-type. **D.** Rea1 depletion affects the association of Rsa4 with pre-60S particles. REA1 (Rea1 +) or GAL::rea1 (Rea1 -) strains expressing HA-tagged Ytm1 or Rsa4 were grown in glucose-containing medium. Immunoprecipitation experiments were then carried out with anti-HA agarose beads and the indicated pre-rRNAs in the input and immunoprecipitated (IP samples) were detected by northern analyses. **E.** Rea1 mutations affect the association of Rsa4 with pre-60S particles. Upper panel: Analysis as in D., except that a GAL::rea1 strain expressing HA-tagged Rsa4 and the indicated Rea1 mutants or wild-type Rea1 (Rea1_wt_) from plasmids was used. EV: strain transformed with an empty vector. Detection of 27SB pre-rRNA by northern hybridization. Lower panel: ratios of the levels of precipitated 27SB pre-rRNA over input levels. **F.** Deleting the α-helical or globular extension or disrupting the D2915-R2976-D3042 salt-bridge network leads to nuclear pre60S particle export defects.

We next investigated whether the pre-rRNA processing defects correlate with a perturbation in the association of Rea1 substrates, Ytm1 and Rsa4, with pre-60S particles. We immunoprecipitated HA-tagged versions of Ytm1 and Rsa4 and analysed the co-immunoprecipitated pre-rRNAs using northern analyses. To our surprise, we observed that Rea1 depletion does not have a major impact on the stability of the association of HA-tagged Ytm1 with 27S pre-rRNAs (Figure 7D). However, Rea1 depletion leads to a significant drop in the co-precipitation efficiency of 27SB pre-rRNA with HA-tagged Rsa4 (Figure 7D). We envisage that in the absence of Rea1, a failure to correctly remodel early Ytm1-containing pre-60S particles perturbs the later association of Rsa4 with incorrectly assembled intermediate pre-60S particles containing the 27SB pre-rRNA. We therefore then used HA-Rsa4 association with 27SB pre-rRNA as a readout of mutant Rea1 activity. In strains expressing Rea1_Δ2916-2974_, Rea1_Δ3072-3244_ or Rea1_D2915A-R2976AD3042A_ the association of HA-Rsa4 with 27SB pre-rRNA was reduced relative to the wild-type control (Figure 7E). In the case of Rea1_Δ2916-2974_ and Rea1_Δ3072-3244_ this reduction was similar to that obtained in the total absence of Rea1, suggesting that these mutants are fully inactive. In contrast, expression of Rea1_D2915A-R2976AD3042A_ had a lesser impact on the association of HA-Rsa4 with 27SB pre-rRNA, consistent with residual activity. Consistent with the pre-rRNA processing and Ytm1/Rsa4 release defects, all Rea1 mutants lead to a major inhibition of pre-60S particle export (Figure 7F).

These results establish the linker middle domain as a critical hub for Rea1 linker remodelling. The α-helical extension is needed to fully sample all the states associated with ATP-hydrolysis suggesting it is indeed involved in communicating ATP-hydrolysis driven conformational changes from the AAA+ ring into the linker. The globular extension of the middle domain functions as a critical structural support for the architecture of the Rea1 linker. The conserved D2915-R2976-D3042 salt-bridge network is required for the correct rotation of the linker top and middle domains around the long linker axis during remodelling. Even though the long linker axis rotation is part of the intrinsic structural flexibility of the Rea1 linker and does not require ATP-hydrolysis, it nevertheless is of functional importance. The rotation is required to ensure the correct orientation of the linker top prior to its ATP-hydrolysis driven engagement with the AAA+ ring. The inability to correctly sample the linker states associated with ATP-hydrolysis correlates with functional defects in pre60S particle maturation and nuclear export.

## Discussion

The results presented here reveal the general principles of the Rea1 mechanism (Figure 8), one of the largest and most complex ribosome maturation factors. We have demonstrated that the Rea1 linker is a functionally important structural element that undergoes an elaborated series of remodelling events. Two general aspects characterize and connect these remodelling events. The first aspect is the swing of the linker top and middle domains towards the AAA+ ring docked MIDAS domain with the region in-between the linker middle and stem domains acting as pivot point. The second aspect is the rotation of the linker top and middle domains around the long linker axis. A particularly interesting feature of this remodelling pathway is the occurrence of events that are part of the intrinsic structural flexibility of the linker like states 1 – 5. Even though these remodelling events do not depend on nucleotide binding or energy provided by ATP-hydrolysis, they still play an important functional role. They ensure that linker top domains are in close proximity to the AAA+ ring and have the correct orientation prior to the ATP-hydrolysis driven engagement with the AAA+ ring. The latter aspect is particularly highlighted by the inability of the Rea1_D2915A-R2976A-D3042A_ mutant to correctly rotate its linker (Figure 6B) and the subsequent functional defects in pre60S particle maturation and nuclear export (Figure 7).

**Figure 8:**
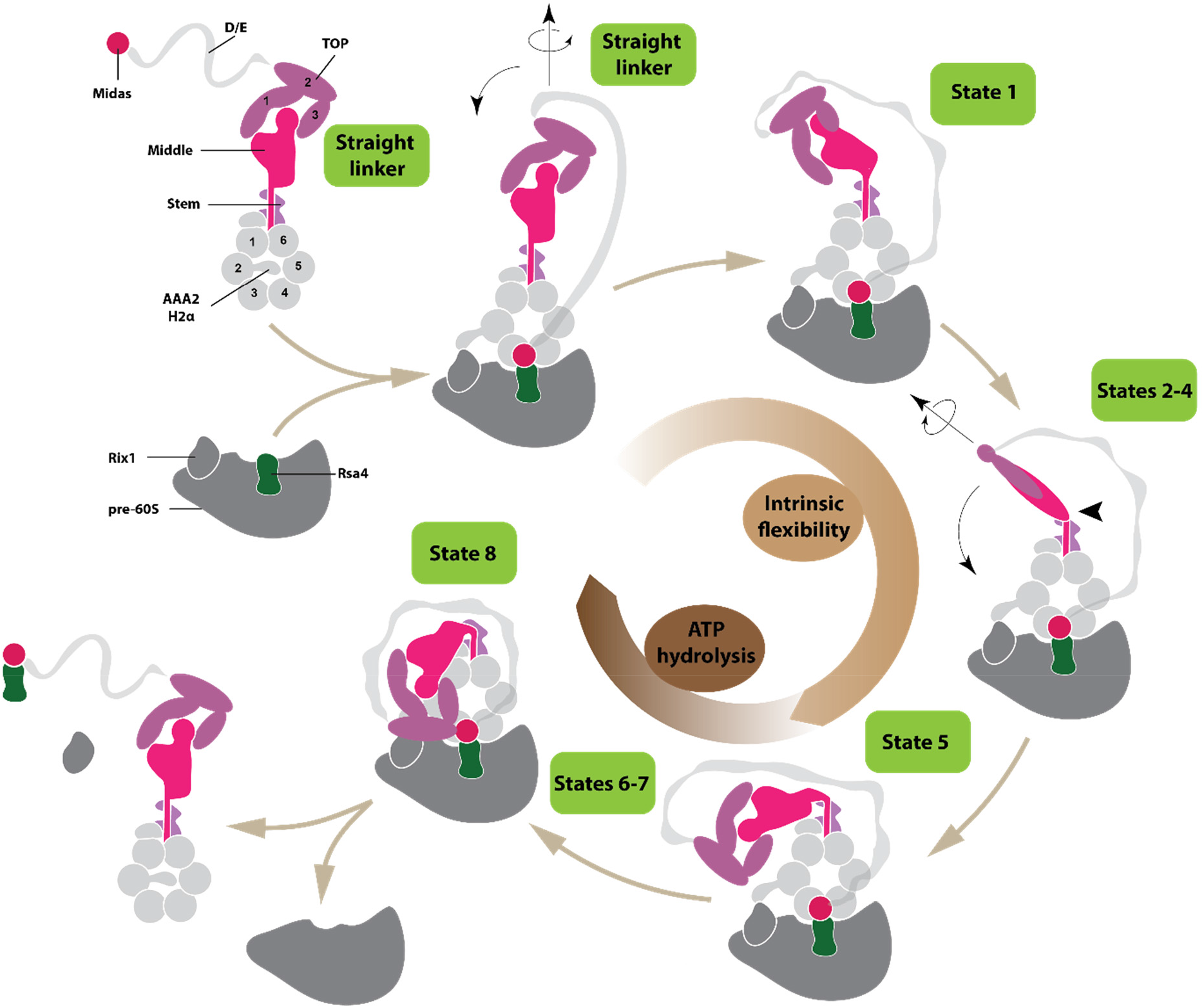
Model for the Rea1 mechanism. Rea1 in the absence of pre60S particles exists in an autoinhibited form with the AAA2H2α insert occupying the central pore of the AAA+ ring and the linker in the straight conformation. The binding of Rea1 to pre60S particles relocates AAA2H2α towards the pre60S particles, which allows the MIDAS domain to dock onto the AAA+ ring to engage with its assembly factor substrate (here Rsa4). The linker remains in the straight conformation. The linker subsequently rotates and pivots towards the plane of the AAA+ ring to reach state 1. From here the linker middle and top domains rotate around the long linker axis and swing towards the AAA+ ring. The region between the linker middle and stem domains acts as pivot point. In state 5, the linker middle and top domains are fully rotated and in close proximity to the AAA+ ring. In states 6-8, the linker top engages with the AAA+ ring and in the final remodelling step state 8 the linker top2 domain interacts with the MIDAS domain to allow the transmission of force for assembly factor removal. Up to state, the linker remodelling is nucleotide independent and driven by the intrinsic conformational flexibility of the linker. States 6 – 8 require ATP hydrolysis.

ATP-hydrolysis is required to engage the rotated linker top and middle domains with the AAA+ ring during states 6 – 8. The ultimate goal of the linker remodelling pathway is to bring the linker top into contact with the AAA+ ring-docked and substrate-engaged MIDAS domain. We have demonstrated that Rea1 linker remodelling is able to produce mechanical force, which would allow the linker top to pull on the substrate engaged MIDAS domain to remove assembly factors form pre60S particles. Since the construct used in the microtubule gliding assays lacked large parts of D/E-rich region and the complete MIDAS domain, these structural elements are dispensable for force production.

We consistently detected that the Rea1 linker states associated with ATP-hydrolysis are sampled only by a minority of particles in our data sets. One possible explanation could be that these states are simply transient and extremely short-lived during the mechanochemical cycle of Rea1. Another possibility is that additional factors are required to further stabilize them such as protein/RNA interaction partners provided in the context of pre60S particle binding.

We have focused our analysis on the linker remodelling states that consistently appeared in comparable negative stain EM data sets. States 1 – 5 suggest that the swing of the linker towards the AAA+ ring and the rotation of the linker top are correlated events, i.e. the linker rotates as it swings towards the AAA+ ring. However, this correlation is not completely strict as in some of the data sets additional linker extended and intermediate states appeared suggesting that the linker can also swing without rotation or fully rotate before reaching the proximity of the AAA+ ring (Supplementary figure 15). The occurrence of these states indicates that alternative pathways exist to remodel the linker from state 1 to state 5.

It is not clear why Rea1 would follow such an elaborate scheme to remove assembly factors from pre60S particles. One possibility is that states 1 – 5 constitute a sensing mechanism. After binding to pre60S particles has occurred, the intrinsic conformational flexibility represented by these states would allow the linker top to come close to the AAA+ ring – pre60S interface to probe if essential maturation events have occurred, before ATP-hydrolysis dependent linker remodelling proceeds to bring the linker top in contact with the AAA+ ring docked MIDAS domain to trigger assembly factor removal.

Currently, there are two mechanistic models for Rea1 functionality. Early structural studies on Rea1 favoured a linker remodelling hypothesis to remove assembly factors from pre60S particles [10], whereas more recently an alternative AAA+ ring model was suggested [19, 22]. In this model, ATP-driven conformational changes in the AAA+ ring are directly communicated to the AAA+ ring docked and substrate-engaged MIDAS domain to produce force for pre60S assembly factor removal. The latter model attributes a more passive role to the Rea1 linker suggesting it acts as a “fishing post” for the flexibly attached MIDAS domain and is not involved in force production [19]. The results presented here favour the linker remodelling hypothesis. Especially the data on the Rea1_Δ2916-2974_ and Rea1_D2915A-R2976A-D3042A_ mutants suggest that a potential AAA+ ring based force production for assembly factor removal is not essential to the Rea1 mechanism. Our structural characterization of these mutants clearly indicates that AAA+ ring and linker are well folded, but they nevertheless show severe functional defects. While these defects can be explained by the linker remodelling hypothesis, the AAA+ ring model would expect these mutants to be functional. However, it is still possible that AAA+ ring based force production plays a supportive role in linker driven assembly factor removal.

Together with the motor protein dynein and the E3 ubiquitin ligase RNF213, Rea1 forms a special subclass within the AAA+ field. All these proteins have their six AAA+ modules concatenated into a single gene, whereas the hexameric AAA+ rings of the vast majority of AAA+ machines are formed by association of individual protomers. In the case of dynein and Rea1, ATP-hydrolysis in the AAA+ ring drives the remodelling of an AAA+ ring linker extension, which in both cases correlates with the production of mechanical force. The dynein linker is with around 400 amino-acid residues much smaller than the Rea1 linker, which spans over around 1700 amino-acid residues, and there are no structural similarities between them. The dynein linker switches between a straight post-powerstroke and bent pre-powerstroke conformation [26] illustrating that the structural heterogeneity is much smaller compared to the Rea1 linker, which is able to adopt at least 9 different conformations (straight linker conformations + states 1 – 8). These differences might be explained by the purpose of force production in dynein and Rea1. The processive movement along microtubules requires highly repetitive linker remodelling. In this context the switch between two remodelling states ensures efficient and fast force production. Rea1 on the other hand functions as a quality control machine. The main task is not highly repetitive force production but rather a single force production event at a precisely controlled time point. In this context many linker remodelling steps offer the opportunity to integrate various signals from the pre60S environment to trigger assembly factor removal only when essential pre60S maturation events have occurred. The linker of the E3 ubiquitin ligase RNF213 is similar to the straight post-powerstroke dynein linker [27]. However, so far no alternative linker remodelling states have been reported for RNF213. It has recently been demonstrated that ATP binding to the AAA+ ring stimulates the ubiquitin ligase activity [28], but it is not clear if this activity increase correlates with RNF213 linker remodelling.

This study highlighted the general mechanistic principles of Rea1. Fully understanding the molecular basis of the Rea1 mechanism will require high resolution investigations on the various linker remodelling states. Especially the AAA+ ring engaged states will be of great interest as these are the ones associated with ATP-hydrolysis and mechanical force production. The extreme low abundance of these states and the structural flexibility of the linker will make such investigations challenging.

## Supporting information

Linker states 1-8 aligned on AAA+ ring

Linker states 1-8 aligned on long linker axis

Linker remodelling is able to create force - example 1

Linker remodelling is able to create force - example 2

Linker remodelling is able to create force - example 3

The linker in the Rea1 D2915A-R2976A-D3042A mutant does not rotate during remodelling

## Acknowledgement

We thank Andrew Carter for providing a plasmid harbouring the SRS-dynein MTBD construct. This work was supported by the collaborative ANR PRC grant “Rea1Com” to H.S. and A.H. This study was further supported by the grant ANR-10-LABX-0030-INRT, a French State fund managed by the Agence Nationale de la Recherche under the frame program Investissements d’Avenir ANR-10-IDEX-0002–02. The authors acknowledge the support and the use of resources of the French Infrastructure for Integrated Structural Biology (FRISBI) ANR-10-INBS-05 and of Instruct-ERIC. We also thank the IGBMC light microscopy (Elvire Guiot, Erwan Grandgirard and Bertrand Vernay) and electron microscopy (Corinne Crucifix and Alexandre Durand) platforms. This work of the Interdisciplinary Thematic Institute IMCBio, as part of the ITI 2021-2028 program of the University of Strasbourg, CNRS and Inserm, was supported by IdEx Unistra (ANR-10-IDEX-0002), and by SFRI-STRAT’US project (ANR 20-SFRI-0012) and EUR IMCBio (ANR-17-EURE-0023) under the framework of the French Investments for the Future Program.

## Material and Methods

### Molecular cloning, protein expression and purification

The Gibson assembly approach was used for all molecular cloning. The centromeric plasmids used for the *S. cerevisiae* pre60S export assay, tetrad dissection as well as the pre-rRNA processing analysis harboured a *ura3* selection marker and the endogenous *rea1* promotor and terminator regions. The Rea1_w_t and mutant constructs were cloned in-between the promotor and terminator. The Rea1Δ_middle-top_ construct was generated by fusing the α-helical extension of the middle domain to the linker stem with a GG spacer. The D/E rich region and the MIDAS domain was then fused to the C-terminal end of the α-helical extension via a GSGS spacer (Rea1Δ_middle-top_: M1-S2608+GG+D2911-D2967+GSGS+F4049-S4910). For the deletion of the α-helical extension of the linker middle domain, Rea1_Δ2916-2974_, residues P2916-V2974 were deleted and replaced with a GGGG spacer. To generate the deletion of the globular extension of the linker middle domain, Rea1_Δ3072-3244_, residues F3072-L3244 were deleted without introducing an additional spacer sequence. The alanine mutations for the Rea1_D2915A-R2976A-D3042A_ salt-bridge network mutant were introduced by mismatched primers.

The recombinant expression and purification of *S. cerevisae* Rea1_wt_, Rea1_ΔAAA2H2α_, Rea1_ΔAAA2H2α+ΔMIDAS_, Rea1_ΔAAA2H2α+Δ4168-4907_, Rea1_Δ2916-2974_, Rea1_Δ3072-3244_ and Rea1_D2915A-R2976A-D3042A_ was done as recently described [17]. The construct Rea1_ΔAAA2H2α+Δ4168-4907_ used in the microtubule gliding assays was further modified by fusing the spytag [25] with two flanking spacer glycines (GAHIVMVDAYKPTKG) between E3943 and T3944 in the linker top2 domain and a GFP with two flanking spacer glycines in the AAA5S domain between S1902 and I1903. Briefly, the constructs were cloned into a pYES2 vector harbouring a Pgal promotor followed by an N-terminal tandem Protein-A tag followed by two preScission protease cleavage sites. The corresponding plasmids were transformed into *S. cerevisiae* strain JD1370 using the Ura selection marker and expression of the constructs was induced by shifting cells to galactose containing media. Cells were pelleted, resuspended in water, flash frozen in liquid nitrogen and stored at −80 ° C. Frozen pellets were blended and further purified by IgG sepharose affinity and gel filtration chromatography as described in detail in [17] with the exception that 50 mM Hepes 7.5, 150 mM NaCl, 2 mM MgCl2 and 1 mM DTT was used as gel filtration buffer.

The spycatcher-dynein MTBD construct was generated by fusing the C-terminus of a chimeric seryl-tRNA synthetase-22:19 dynein MTBD construct in the high microtubule affinity state [29] to the to the N-terminus of the spycatcher domain [25] in a pET42a vector background. The construct was expressed in *Escherichia coli* strain BL21(DE3) by induction with 1mM isopropyl β-D-thiogalactopyranoside for 16 h at 18 ° C. All subsequent steps were carried out at 4 ° C. The cells were harvested by centrifugation at 5000 rpm for 15 min. The pellet was resuspended in lysis buffer (50 Hepes 7.5, 200 mM NaCl, 10% glycerol, 1mM 2-mercaptoethanol, 5 mM imidazole, 0.5 mM phenylmethylsulfonyl fluoride) and lysed using a sonicator. The lysate was subsequently clarified by centrifugation at 30000 rpm for 30 min. The expressed protein was incubated with nickel-nitrilotriacetic acid beads for 1h and washed with three column volumes of wash buffer (50 Hepes 7.5, 200 mM NaCl, 10% glycerol, 1mM 2-mercaptoethanol, 30 mM imidazole, 1mM Mg-ATP). The protein was eluted with elution buffer (50 Hepes 7.5, 200 mM NaCl, 10% glycerol, 1mM 2-mercaptoethanol, 250 mM imidazole).

### Yeast strains and media

For the pre60S export assay, an auxotrophic *S. cerevisiae* strain (-ura, -leu, -his, -trp) featuring an mCherry fused to the C-terminus of the endogenous histone 2B locus was used. The strain was further modified by tagging the C-terminus of the endogenous Rea1 with the IAA17 protein using kanamycin selection and IAA17 partner protein OsTir1 was integrated into the H0 locus using hygromycin selection. The addition of auxin to the medium triggers the IAA17-OsTir1 interaction and subsequent proteasome degradation [30]. This strain was used to transform a pRS415 plasmid harbouring Rpl25-GFP using the -leu selection marker. The strain was subsequently modified by transforming the centromeric plasmids harbouring Rea1_w_t, Rea1Δ_middle-top_, Rea1_Δ2916-2974_, Rea1_Δ3072-3244_ and Rea1_D2915A-R2976A-D3042A_ constructs (see below) using -leu -ura double selection.

To construct the diploid REA1/rea1::kan^R^ strain in the BMA64 genetic background (*MATa/MATα,MATa; his3-11_15/his3-11_15; leu2-3_112/leu2-3_112; ura3-1/ura3-1; trp1Δ2/trp1Δ2; ade2-1/ade2-1; can1-100/can1-100*) for the tetrad dissection assays, the rea1::kan^R^ deletion cassette was mobilized by PCR amplification of genomic DNA extracted from the commercial REA1/rea1::kan^R^ diploid strain in the BY4741 background (purchased from Open Biosystems), using primers OHA532 and OHA534.

GAL::REA1 strain was obtained by transforming haploid BY4741 strain (*MATa*, *his3Δ1*, *leu2Δ0*, *met15Δ0*, *ura3Δ0*) with a kan^R^-PGAL1 cassette flanked by REA1 sequences corresponding to the promoter region and the beginning of the open reading frame at the 5’ and 3’ ends, respectively. This cassette was produced by PCR amplification of plasmid pFA6a-kanMX6-PGAL1 [31] using oligonucleotides OHA529 and OHA530. Genomic insertion of the cassette has been verified by PCR amplification of genomic DNA purified from selected transformants and oligonucleotides OHA532 and OHA533.

YTM1-HA and GAL::REA1/YTM1-HA strains were obtained by transforming the BY4741 or the GAL::REA1 strains with a 3HA-His3MX6 PCR cassette flanked by YTM1 open reading frame and terminator sequences at the 5’ and 3’ ends, respectively, produced by amplification of plasmid pFA6a-3HA-His3MX6 [31] with oligonucleotides OHA543 and OHA544. Genomic insertion of the cassette has been verified by PCR amplification of genomic DNA purified from selected transformants and oligonucleotides OHA545 and OHA546.

HA-RSA4 and GAL::REA1/HA-RSA4 strains were obtained by CRISPR-Cas9-mediated genome editing of the BY4741 or the GAL::REA1 strains using the procedure described in [32]. Oligonucleotides OHA553 and OHA554 were annealed and cloned into pML104 vector to generate a plasmid expressing the Cas9 nuclease and a guide RNA targeting the nuclease in the vicinity of the ATG of RSA4 open reading frame. This plasmid was transformed into the BY4741 or the GAL::REA1 strains along with a repair PCR fragment consisting in the 3HA tag sequence flanked by RSA4 sequences corresponding to the promoter region and the beginning of the open reading frame at the 5’ and 3’ ends, respectively. This cassette was produced by PCR amplification of plasmid pFA6a-3HA-kanMX6 [31] with oligonucleotides OHA551 and OHA552. Transformants were selected on synthetic medium (see below) lacking uracil to select clones having internalized the recombinant pML104 plasmid. The plasmid was then rapidly counter-selected on synthetic YNB medium containing 5-Fluoroorotic acid. Genomic insertion of the cassette in the resulting clones has been verified by PCR amplification of genomic DNA purified from selected transformants and oligonucleotides OHA559 and OHA560.

Transformed REA1/rea1::kanR diploid strains were sporulated on 2 % potassium acetate, 0.22 % yeast extract, 0.05 % glucose, 0.079 % complete supplement mixture (MP biomedicals), 2 % agar, adjusted to pH 7.0 with potassium hydroxide. Tetrads were dissected using a Singer MSM dissection microscope.

Strains were grown either in YP medium (1 % yeast extract, 1 % peptone) (Becton-Dickinson) supplemented with 2 % glucose or 2 % galactose as the carbon source or in synthetic medium (0.17 % yeast nitrogen base (MP Biomedicals), 0.5 % (NH4)2SO4) supplemented with 2 % glucose or 2 % galactose and the required amino acids. Selection of the kanamycin-resistant transformants was done by addition of G418 to a final concentration of 0.2 mg/ml.

### S. cerevisiae pre60S export assays

Strains carrying the Rpl25-GFP and Rea1 construct plasmids were used to inoculate a 50 ml pre-culture of CSM -leu -ura minimal media enriched with 2% glucose and 0.1 mM Adenine. The pre-culture was incubated over night at 200 rpm and 30 ° C. The following morning the preculture was used to inoculate another 50 ml of CSM -leu -ura minimal media enriched with 2% glucose and 0.1 mM Adenine to a final OD of 0.2. In order to induce the degradation of endogenous Rea1, auxin to a final concentration of 1 mM was added. The culture was incubated at 200 rpm and 30 ° C and cells were imaged after 6h.

### Negative stain electron microscopy

Standard plain carbon grids (copper 300 mesh, vendor: EMS) were glow discharged in air plasma (30s, ~2.5mA, 0.21mbar). Rea1 samples were diluted in 50 mM Hepes 7.5, 150 mM NaCl, 2 mM MgCl2 and 1 mM DTT to a final concentration of 25 nM in the presence of 3mM nucleotide or without nucleotide. 5 μl of the 25 nM sample were applied to the glow discharged grid for 2 min. After side blotting out the excess of sample, grids were stained following standard procedures with freshly centrifuged and filtered 2% Uranium Acetate.

Grids were imaged using a FEI Technai G2 FEG TEM operated at 200kV HT equipped with a charge-coupled device camera (2K US10001; Gatan). Acquisitions were made with SerialEM [33] at 50kx magnification(2.002A/pix), a nominal defocus of −1.5μm and a 5×5 montage mode for speeding up data collection. Micrographs were extracted from montage stacks using IMOD [34] and processed using Relion [35].

The Rea1_ΔAAA2H2α_ ATPγS data set (Supplementary figure 16) consisted of a total of 74141 particles, which was sufficient to resolve all 8 linker remodelling states. The Rea1_wt_ ATP, APO, AMPPNP and ATPγS data sets had 70952, 69908, 76407 and 87826 particles. The Rea1_ΔAAA2H2α_ ATP, ADP, AMPPNP, ATPγS + PhoX and APO data sets consisted of 125294, 97479, 93919, 85123 and 314443 particles. The Rea1_Δ2916-2974_, Rea1_Δ3072-3244_ and Rea1_D2915A-R2976A-D3042A_ ATP data sets consisted of 71448, 140277 and 69331 particles.

### CryoEM electron microscopy

For the Rea1_ΔAAA2H2α_ ATPγS and Rea1_D2915A-R2976A-D3042A_ ATP data sets, standard copper/rhodium quantifoil R2/2 with an additional carbon layer and streptavidin affinity grids [36] were used. In addition, standard copper/rhodium quantifoil R2/2 without additional carbon layer were used for Rea1_D2915A-R2976A-D3042A_ ATP. 3 μl of Rea1_ΔAAA2H2α_ or Rea1_D2915A-R2976A-D3042A_ at a concentration of 150 nM in 50 mM Hepes 7.5, 150 mM NaCl, 2 mM MgCl_2_, 1 mM DTT, 0.005% DDM and 3 mM ATPγS or ATP were incubated with plasma cleaned (80% oxygen / 20% argon, 30% power, 90 sec) carbon grids. In case of the streptavidin affinity grids, 3 μl of biotinylated Rea1_ΔAAA2H2α_ or Rea1_D2915A-R2976A-D3042A_ at a concentration of 60 nM were used. The streptavidin affinity grids were further treated as described recently [36]. For the unsupported Rea1D2915A-R2976A-D3042A ATP grids, 3 μl at a concentration of 1800 nM in the buffer described above were used. All grids were plunge frozen with a Vitrobot Mark IV robot (FEI), maintained at 95% humidity and 10 °C.

All data sets were collected on a FEI Titan Krios operating at 300 kV equipped with a Cs corrector and a Gatan K3-Summit detector using a slit width of 20 eV on a GIF-quantum energy filter (Gatan), except for the streptavidin affinity dataset, which was collected on a Thermo Fisher Glacios operating at 200 kV equipped with a K2-Summit direct electron detector (Gatan). In case of the carbon supported grids, two data sets where acquired at 0° and 30° tilt. All Krios datasets were acquired at 81kx magnification (0.862 A/pix). The Glacios datasets were acquired at 45kx magnification (0.901 A/pix). Acquisitions were performed semi-automatically, managed with Serial EM [33].

Micrographs were processed with relion [35]. The streptavidin crystal pattern in the micrographs of the streptavidin affinity grids was erased in Fourier space using python scripts described in [36]. Particles were extracted with a binning of 4 times. The combined particles datasets were submitted to 2D classifications, stochastic gradient decent to create initial models and several rounds of 3D classification from which the straight linker conformation emerged as the only interpretable cryoEM map in the case of the Rea1_ΔAAA2H2α_ ATPγS data set. For the Rea1_D2915A-R2976A-D3042A_ ATP data set, three linker conformations (I-III) could be obtained with conformation I being highly similar to the straight linker conformation of Rea1_ΔAAA2H2α_ ATPγS cryoEM map. The Rea1_ΔAAA2H2α_ ATPγS data set had 230k particles. The final map of the straight linker as shown in Figure 3A consisted of 100k particles and refined to Nyquist spacing (7.1 Å for a pixel size of 3.45 Å/pixel). The Rea1_D2915A-R2976A-D3042A_ data set had 1035k particles. The final models for Rea1_D2915A-R2976A-D3042A_ ATP conformations I – III consisted of 300k, 250k and 240k particles and also refined to Nyquist spacing (≈ 7 Å for a pixel size 3.45 Å/pixel). The Rea1_ΔAAA2H2α_ ATPγS and Rea1_D2915A-R2976A-D3042A_ ATP (conformation I, II, III) maps have been submitted to the electron microscopy data bank with access codes EMD-XXXX, EMB-XXXX, EMD-XXXX and EMD-XXXX.

### Crosslinking mass spectrometry

#### Sample preparation and PhoX crosslinking

Rea1_ΔAAA2H2α_ in 50mM Hepes, 150mM NaCl, 2mM MgCl2 was incubated with 5mM ATPγS or 5mM AMP-PNP. The final protein concentration after nucleotide incubation was 1.25 mg/ml. A solution of bovine serum albumin was prepared at 2 mg/ml (in PBS) as a quality control sample of the crosslinking reaction. A 2 mg aliquot of PhoX crosslinker (Disuccinimidyl Phenyl Phosphonic Acid, Bruker) [37] was freshly diluted in DMSO to a final concentration of 5.16 mM. Each Rea1_ΔAAA2H2α_ sample was split into three aliquots (3x ATPγS, 3x AMPPNP) of 15 μg protein each and subsequently incubated with 1μL of PhoX crosslinker stock solution, corresponding to a 200 molar excess of PhoX. The quality control Bovine Serum Albumin triplicate was also crosslinked with a 100 molar excess of PhoX. The crosslinking reaction was carried out at 20°C for 45 min and quenched by adding Tris-HCl to a final concentration of 10mM and an additional 20 min incubation step. A fourth 15 μg ATPγS aliquot was crosslinked and kept for negative-stain electron microscopy analysis (Supplementary figure 9).

To control the reaction efficiency, 1.4 μg of the non-crosslinker and crosslinked samples were kept for a mass photometry analysis to check for the appearance of non-specifically crosslinked dimers (Supplementary figure 9). After the quality check, all samples were reduced by adding DTT to a final concentration of 5 mM and incubation at 37°C for 30 min. The subsequent alkylation was carried out by adding Iodoacetamide to a final concentration of 15 mM followed by an 1 hour incubation step in the dark. The samples were further processed by digesting them overnight with a Trypsin/Lys-C mix (Promega, Madison, USA) at a 50:1 substrate:enzyme ratio (w/w) at 37°C overnight. The digestions were finally quenched with 1% TFA.

#### Automatized peptide cleanup and enrichment of PhoX-crosslinked peptides

Peptides were first cleaned up by using the AssayMAP Bravo platform (Agilent Technologies; Santa Clara, California) with 5μL C18 cartridges (Agilent). Cartridges were primed with 100 μl 0.1% TFA in 80% ACN and equilibrated with 50 μl 0.1% TFA in H2O. 180 μl of digested peptides diluted in equilibration buffer were loaded on the cartridges and washed with 50 μl equilibration buffer. Peptides were eluted with 50 μl 0.1% TFA in 80% ACN. In order to enrich the PhoX-crosslinked peptides, 5 μl Fe(III)-NTA cartridges (Agilent) were primed with 100 μL of 0.1% TFA in H2O and equilibrated with 50 μl 0.1% TFA in 80% ACN [37]. 140 μl of the C18-eluted peptides diluted in 0.1% TFA in 80% ACN were loaded and subsequently washed with 50 μl 0.1% TFA in 80% ACN. PhoX-crosslinked peptides were eluted with 50 μl of 1% NH4OH solution and stored at −80°C prior to the mass spectrometry analysis.

#### LC-MS/MS acquisition and data processing

Before being injected, PhoX-crosslinked peptides were dried in a SpeedVac concentrator and resuspended in 10 μl of 2% ACN/0.1% formic acid. NanoLC-MS/MS analysis was performed using a nanoAcquity Ultra-Performance-LC (Waters, Milford, USA) hyphenated to a Q-Exactive Plus Orbitrap mass spectrometer (Thermo Fisher Scientific, Bremen, Germany) equipped with a nanoSpray source. Samples were first trapped on a nanoACQUITY UPLC precolumn (C18, 180 μm × 20 mm, 5 μm particle size), prior to separation on a nanoACQUITY UPLC BEH130 column (C18, 75 μm × 250 mm with 1.7 μm particle size, Waters, Milford, USA) maintained at 60°C. A Gradient of 105 min was applied using mobile phases A (0.1% v/v formic acid in H2O) and B (0.1% v/v formic acid in ACN). The following conditions were applied: 1–3 % B for 2 min, 3–35% B for 77 min, 35–90% B for 1 min, 90% B for 5 min, 90–1% B for 2 min and finally 1% B maintained for 2 min (flow rate of 400 nl/min). The Q Exactive Plus Orbitrap source temperature was set to 250°C and the spray voltage to 1.8 kV. Full scan MS spectra (300–1,800 m/z) were acquired in positive mode (resolution of 140,000, max. injection time of 50 ms, AGC target value of 3.106) with the lock-mass option enabled (polysiloxane ion from ambient air at 445.12 m/z). Using a Data Dependant Acquisition strategy, the 10 most intense peptides per full scan (charge states > 2) were isolated using a 2 m/z window and fragmented using stepped collision energy HCD (27%, 30%, 33% normalized collision energy). MS/MS spectra were acquired with a resolution of 35 000, a maximum injection time of 100 ms, an AGC target value of 1.105, and a dynamic exclusion time of 60 s. NanoLC-MS/MS system was piloted with XCalibur software v3.0.63, 2013 (Thermo Scientific) and a NanoACQUITY UPLC console v1.51.3347 (Waters).

Raw data were directly processed with Thermo Proteome Discoverer 2.5.0.400 (Thermo Scientific) using the XlinkX [38] node for identification of crosslinks and the Sequest HT node for the identification of linear peptides. For both linear and crosslinked peptides searches, Cystein carbamidomethylation was set as fixed modification. Methionine oxidation, N-term acetylation, tris-quenched monolinks and water-quenched monolinks were set as dynamic modifications. Trypsin was set as the cleavage enzymes with minimal length of 7 amino acids, 2 (linear peptides) and 3 (crosslinked peptides) missed cleavages were allowed. To increase confidence, identification were only accepted for a minimal score of 40 and a minimal delta score of 4 [37]. A 1% false discovery rate was also applied at a crosslinked peptides level (XlinkX validator node). In-house database for linear peptides identification was composed of 187 entries (Rea1_ΔAAA2H2α_, common contaminants and reversed sequences), database for crosslinks identification was only composed of the Rea1_ΔAAA2H2α_ sequence (purified sample) to reduce the search space. For the consensus step, proteins identifications were controlled at a 1% FDR in the protein validator node. Out of the three replicates performed, cross-linking interactions were validated when seen in at least 2 out of 3 replicates. The XL-MS dataset has been deposited to the ProteomeXchange Consortium via the PRIDE51 partner repository with the dataset identifier PXDxxxxx.

#### Mass Photometry

Mass Photometry (TWOMP, Refeyn Ltd, Oxford, UK) was performed on the crosslinked ATPγS and AMPPNP Rea1_ΔAAA2H2α_ replicates as well as on the ATPγS Rea1_ΔAAA2H2α_ non-crosslinked control. Microscope cover slides (24×50mm, 170±5μm, No. 1.5H, Paul Marienfeld GmbH & Co. KG, Germany) were cleaned with milli-Q water, isopropanol, milli-Q water and were then dried with a clean nitrogen stream [39]. Six-well reusable silicone gaskets (CultureWellTM, 50-3mm DIA x 1mm Depth, 3-10 μL, Grace Bio-Labs, Inc., Oregon, USA) were carefully cut and assembled on the cover slide center. After being placed in the mass photometer and before each acquisition, an 18 μL droplet of Phosphate Buffer Saline (PBS) was put in a well to enable focusing on the glass surface. All samples were first diluted with their native buffer, and then 2 μL of the protein stock solution was diluted into the 18 μL PBS droplet. A contrast-to-mass calibration was performed with Bovine Serum Albumin, Bevacizumab, and L-Glutamate Dehydrogenase diluted in PBS buffer, pH 7. For analysis, samples were diluted just below the saturation to obtain the highest number of counts without losing signal quality (typically between 10 to 40 nM in the droplet). Movies of 60 s were recorded in triplicate for all samples with AcquireMP, and processed with DiscoverMP softwares (Refeyn Ltd, Oxford, UK).

### Microtubule gliding assays

90 μl of Rea1_ΔAAA2H2α+Δ4168-4907_ (0.7 mg/ml) were mixed 60 μl of spycatcher-dynein MTBD (3.8 mg/ml) and 150 μl 200 mM NaOH-Pipes pH 6.8 and incubated for 4 h at room temperature to catalyze the formation of the covalent spycatcher-spytag bond. The reaction mixture was subsequently run over a superpose 6 gel filtration column to separate the Rea1_ΔAAA2H2α+Δ4168-4907_ -spycatcher-dynein MTBD construct from the unreacted Rea1_ΔAAA2H2α+Δ4168-4907_ and spycatcher-dynein MTBD samples.

To generate microtubules for gliding assays, 2 μl of porcine tubulin (5mg/ml) were mixed with 0.25 μl of TMR labelled tubulin (5mg/ml, in general tubulin buffer, vendor: Cytoskeleton) and incubated for 5 min on ice. After the addition of 2.5 μl general tubulin buffer, 2.5 μl tubulin glycerol buffer and 0.75 μl 20 mM GTP (Cytoskeleton), the mixture was incubated at 37 ° C for 20 min. To create the microtubule stock solution for the gliding assays, 25 μl of general tubulin buffer enriched with 20 μM taxol (cytoskeleton) were added.

The flow chamber was incubated with 10 μl anti-GFP (Roche, 500 μg/ml in PBS buffer) for 2 min and subsequently washed twice with 10 μl BRB80-Casein (80 mM NaOH Pipes pH 6.8, 2 mM MgCl_2_, 1 mM EGTA, 1 mM DTT and 2 mg/ml Casein). Rea1_ΔAAA2H2α+Δ4168-4907_- spycatcher-dynein MTBD after gel filtration (0.15 mg/ml) was diluted 1:10 in BRB80 and 3 x 10 μl were applied to the flow chamber for 2 min followed by two 10 μl BRB80-Casein washing steps. The motility solution was prepared by adding 1 μl microtubule stock solution and 1 μl gloxy solution (catalase and glucose oxidase in BRB80 buffer) to 98 μl BRB80-Casein, 3 mM Mg-ATP, 40 mM KAc, 20 μM taxol, 0.5% glucose. 10 μl of the motility solution was applied to the flow chamber and after a 2 min incubation step the cover slide was imaged by TIRF microscopy. Slow, directed microtubule movements were occasionally observed in the sample. Such events were not detected with motility solutions without Mg-ATP.

### ATPase assays

For all ATPase assays, the EnzChek Phosphate Assay Kit (Molecular Probes) was used according to the supplier recommendations. The reaction volume was 150 μl consisting of 30 μl 5x assay buffer (150 mM HEPES-NaOH pH 7.2, 10 mM Mg-Acetate, 5 mM EGTA, 50% Glycerol, 5 mM DTT), 30 μl MESG (EnzCheck substrate), 1.5 ml PNP (purine nucleoside phosphorylase), 10 μl Mg-ATP (15 mM) and 30 – 50 nM protein. Experiments were carried out on aGENios spectrophotometer (TECAN). All assays were done in triplicate. The ATPase rates were 0.96 ± 0.17, 0.41 ± 0.03, and 0.54 ± 0.1 mol Phosphate/mol Rea1/s for Rea1_wt_, Rea1_Δ2916-2974_, and Rea1_D2915A-R2976A-D3042A_.

### Miscellaneous

RNA extractions and northern blotting experiments were performed as described [40] and the Anti-HA immunoprecipitation experiments were carried out as described in [41].

