## Supplementary-figures-and-legends for "The Ribosome Maturation Factor Rea1 utilizes nucleotide independent and ATP-hydrolysis driven Linker remodelling for the removal of ribosome assembly factors"

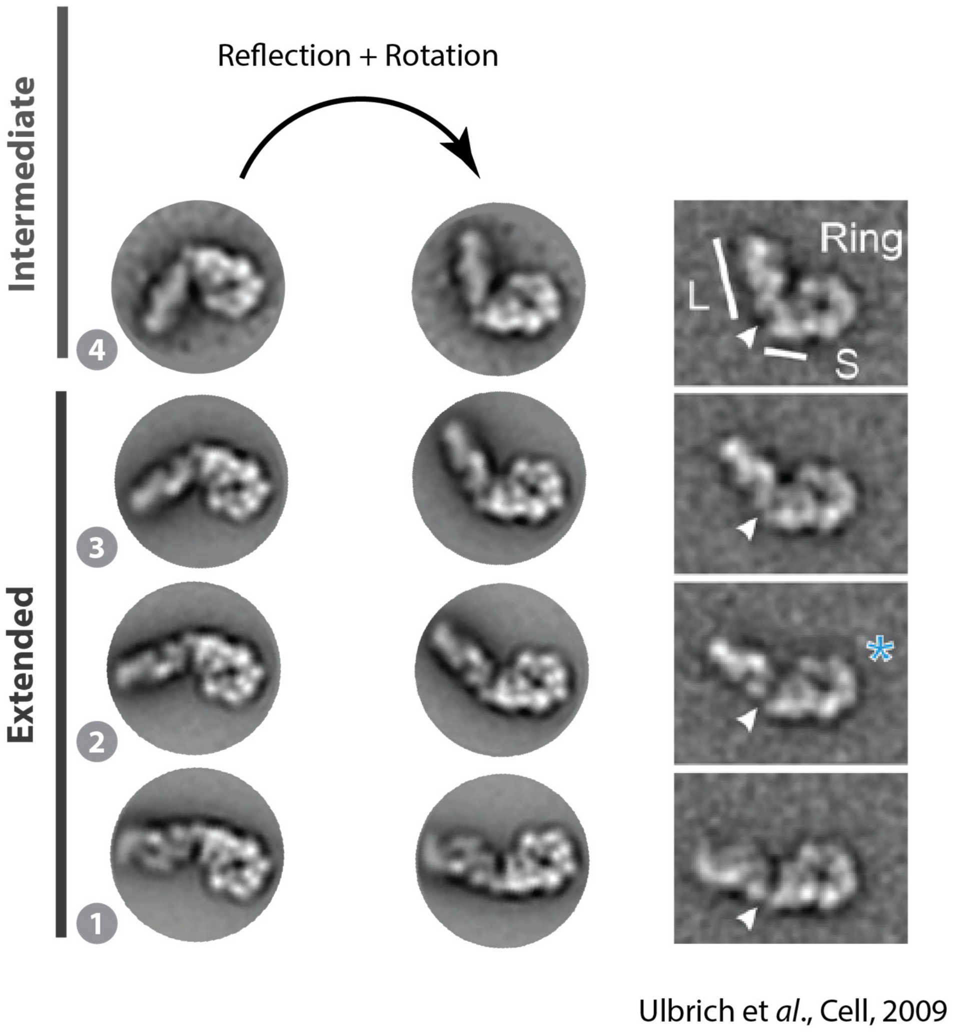


**Supplementary figure 1: Comparison of Rea1$_{wt}$ negative stain EM 2D classes with published data.** States 1 -4 are similar to 2D classes published by Ulbrich et al.

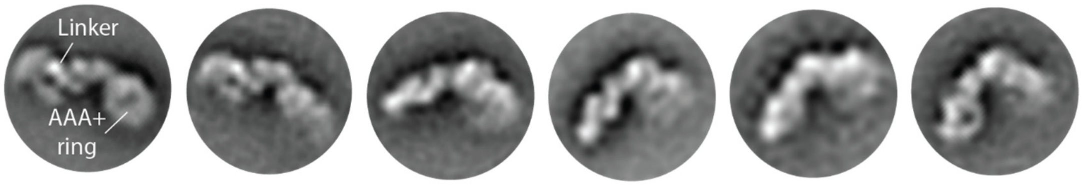


**Supplementary figure 2: Negative stain 2D class averages of Rea1$_{\Delta AAA2H2\alpha}$ in the absence of nucleotide.** Linker conformations consistent with states 1 – 5 of the extended and intermediate classes can be observed.

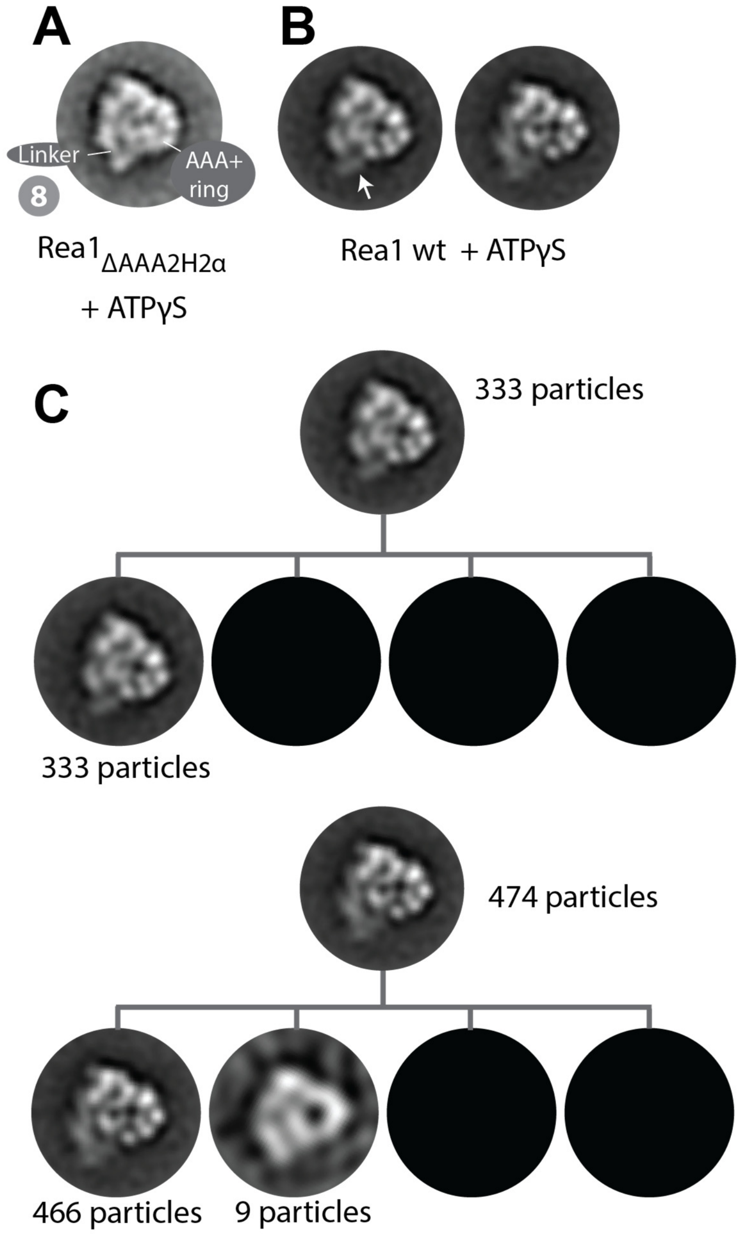


**Supplementary figure 3: Linker remodelling state 8 is not stable in Rea1$_{wt}$. A.** State 8 as observed in Rea1$_{\Delta AAA2H2\alpha}$ in the presence of ATPγS. **B.** Two 2D class averages obtained from a Rea1$_{wt}$ ATPγS data set. The two 2D classes are similar to state 8 of Rea1$_{\Delta AAA2H2\alpha}$, but thin stain for the linker tip (left, white arrow) or the complete linker (right) indicates increased structural flexibility. **C.** The increased structural flexibility of the linker in B. might be due to a mixture of well-folded state 8 particles and partially or completely unfolded particles. To rule out this possibility, we re-classified the particles in B. into 4 sub-classes. The sub-classification brings back the original 2D classes confirming that the linker in these classes is too flexible to stably sample state 8.

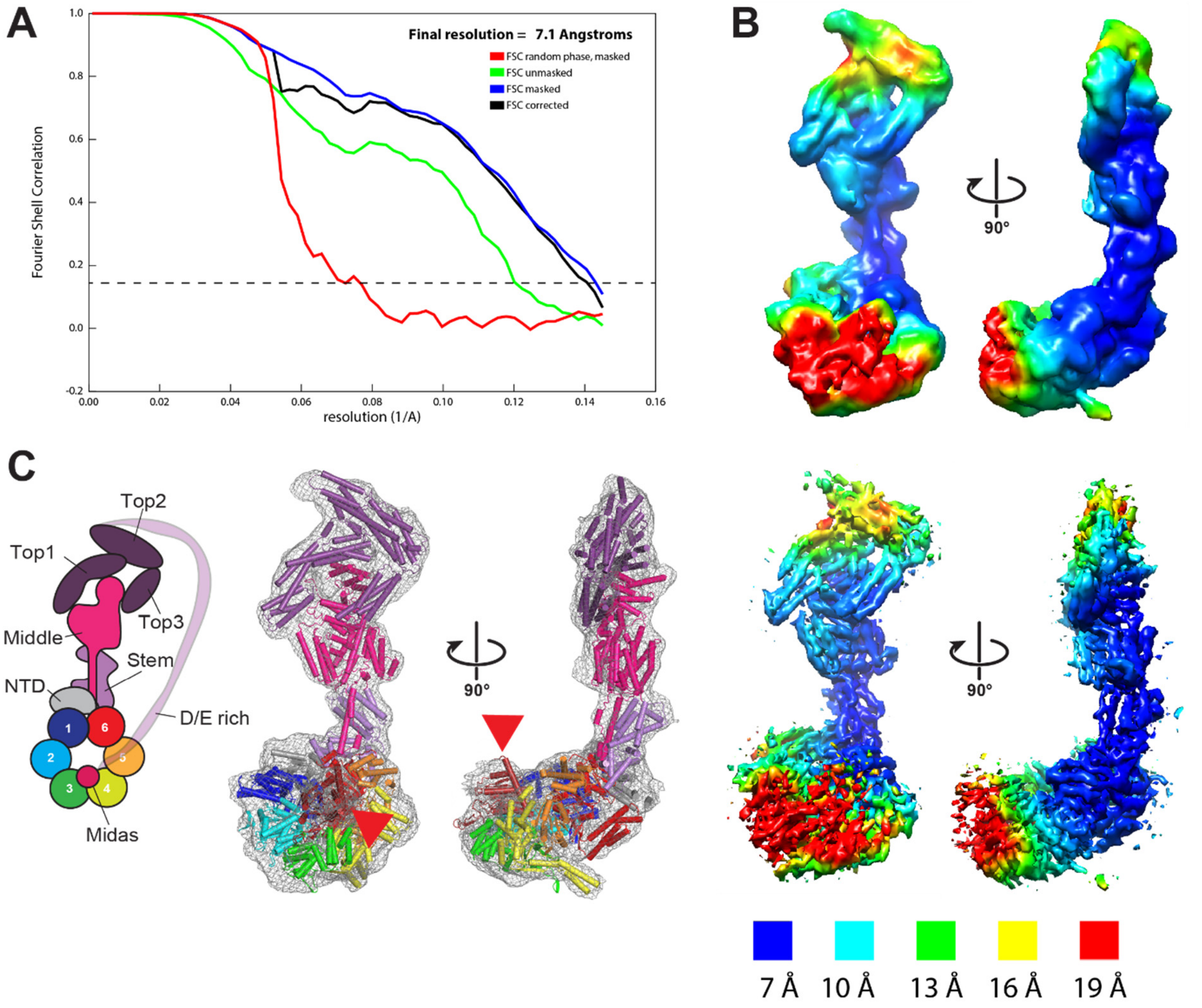


**Supplementary figure 4: Overall quality of the Rea1$_{\Delta AAA2H2\alpha}$ ATPγS cryoEM map. A.** Fourier shell correlation (FSC) plot for half-maps of the 3D reconstruction. The 0.143 FSC criteria is indicated as horizontal dashed line. The final overall resolution is 7.1 Å. **B.** Local resolution map, upper panels: unsharpened map, lower panels: B-factor sharpened map. Secondary structure elements can be identified. **C.** Schematic cartoon representation of structure (left) and match of structure in map (right). Red arrow heads highlight the AAA+ ring docked MIDAS domain.

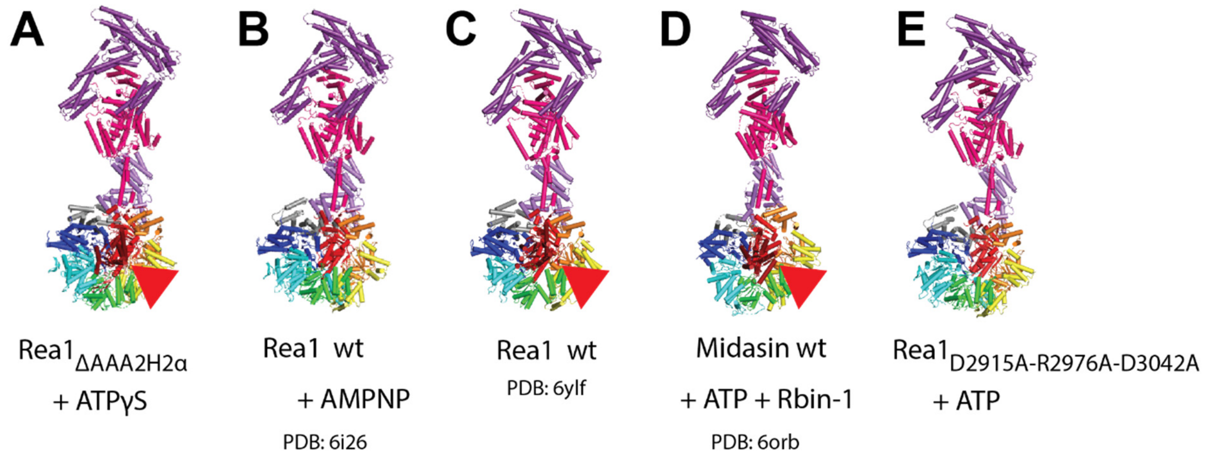


**Supplementary figure 5: Comparison of the straight linker in the Rea1$_{\Delta AAA2H2\alpha}$ ATPγS cryoEM structure with other Rea1/Midasin cryoEM structures. A.** Straight linker in Rea1$_{\Delta AAA2H2\alpha}$ ATPγS structure (this study). **B.** Rea1$_{wt}$ in the presence of AMPNP (Sosnowski et al., 2018). **C.** Rea1$_{wt}$ bound to a pre60S particle (Kater et al., 2020). **D.** Midasin$_{wt}$ in the presence of ATP and the Rea1/Midasin inhibitor Rbin-1 (Chen at al., 2018). **E.** Straight linker in Rea1$_{D2915A\text{-}R2976A\text{-}D3042A}$ (Conformation I.) in the presence of ATP (this study). In all structures the linker adopts a similar straight conformation with respect to the AAA+ ring. Red arrow heads highlight the AAA+ docked MIDAS domain.

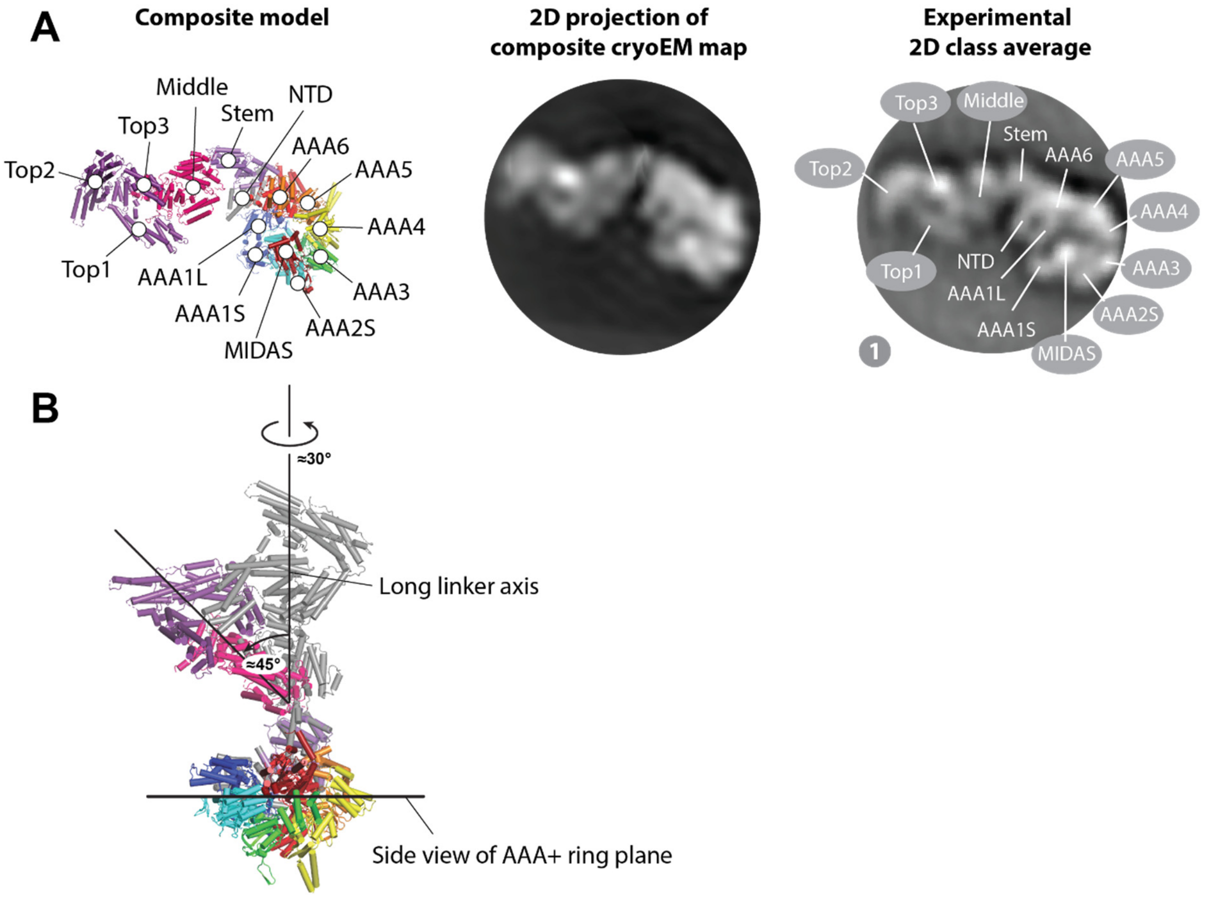


**Supplementary figure 6: The linker in state 1 has moved with respect to the AAA+ ring compared to the straight linker conformation in Rea1$_{\Delta AAA2H2\alpha}$ ATPγS cryoEM structure. A.** Combining the AAA+ ring map in the orientation used to assign the AAA+ sub-domains in state 1 with the linker map in the orientation used to assign linker sub-domains in state 1 by overlapping the linker stem area allowed us to create structural composite model for state 1. **B.** Aligning the AAA+ ring of this composite model with the AAA+ ring in the Rea1$_{\Delta AAA2H2\alpha}$ ATPγS cryoEM structure reveals that the linker top and middle domains have rotated by ≈ 30˚ and swung by ≈ 45˚ towards the AAA+ ring plane. Straight linker of the Rea1$_{\Delta AAA2H2\alpha}$ ATPγS cryoEM structure is shown in grey.

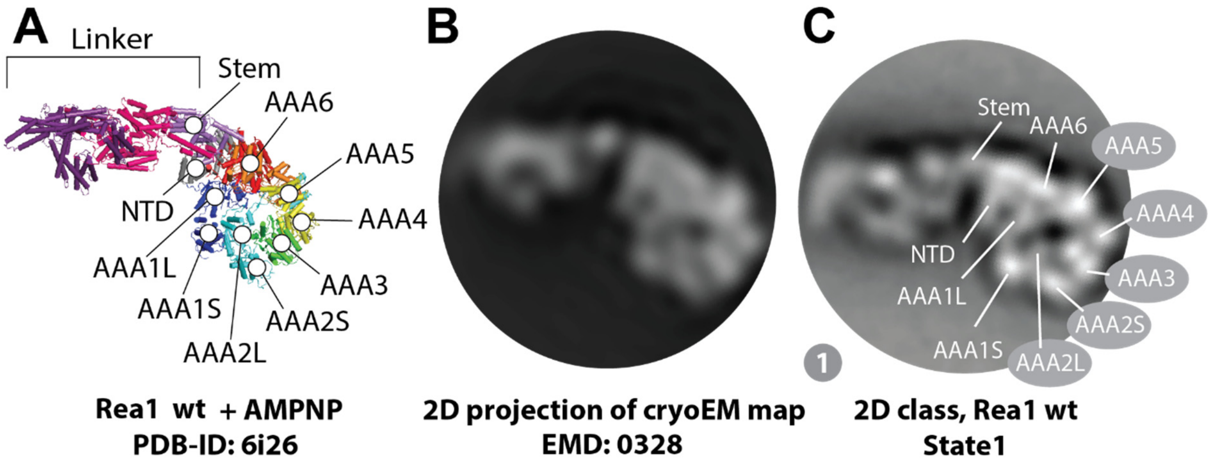


**Supplementary figure 7: The AAA+ ring in the 2D class averages of Rea1$_{wt}$ does not harbour a docked MIDAS domain. A.** Structure of Rea1$_{wt}$ in the presence of AMPPNP (Sosnowski et al., 2018). The AAA+ ring does not feature a docked MIDAS domain **B.** 2D projection of the corresponding cryoEM map of A. low pass filtered to 25 Å. **C.** The AAA+ ring in the 2D class averages of Rea1$_{wt}$ (here state 1 as example) matches well with the projection in B. suggesting the MIDAS domain is not docked onto the AAA+ ring.

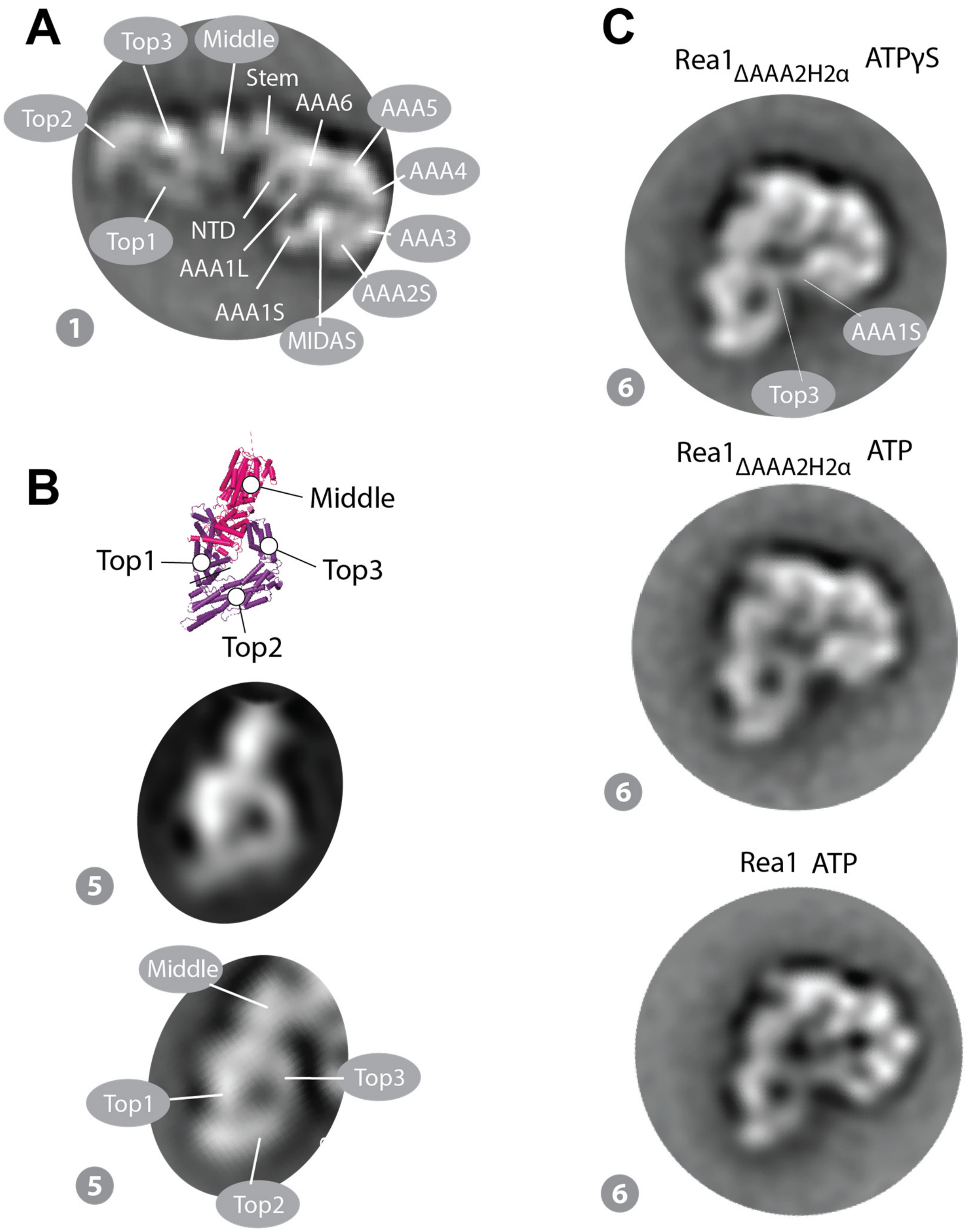


**Supplementary figure 8: State 6 of the AAA+ ring engaged linker conformations features a connection between AAA1S and the linker top3 domain. A.** Domain assignments in the AAA+ ring and linker. **B.** Two upper panels: Linker middle and top domains rotated into state 5 and corresponding cryoEM map projection (compare also Figure 4C and D). Lower panel: Assignment of linker domains in state 5 based on the two upper panels. **C.** The domain assignments in A. and B. suggest a connection between AAA1S and the linker top3 domain in state 6. The connection is visible in three independently collected data sets.

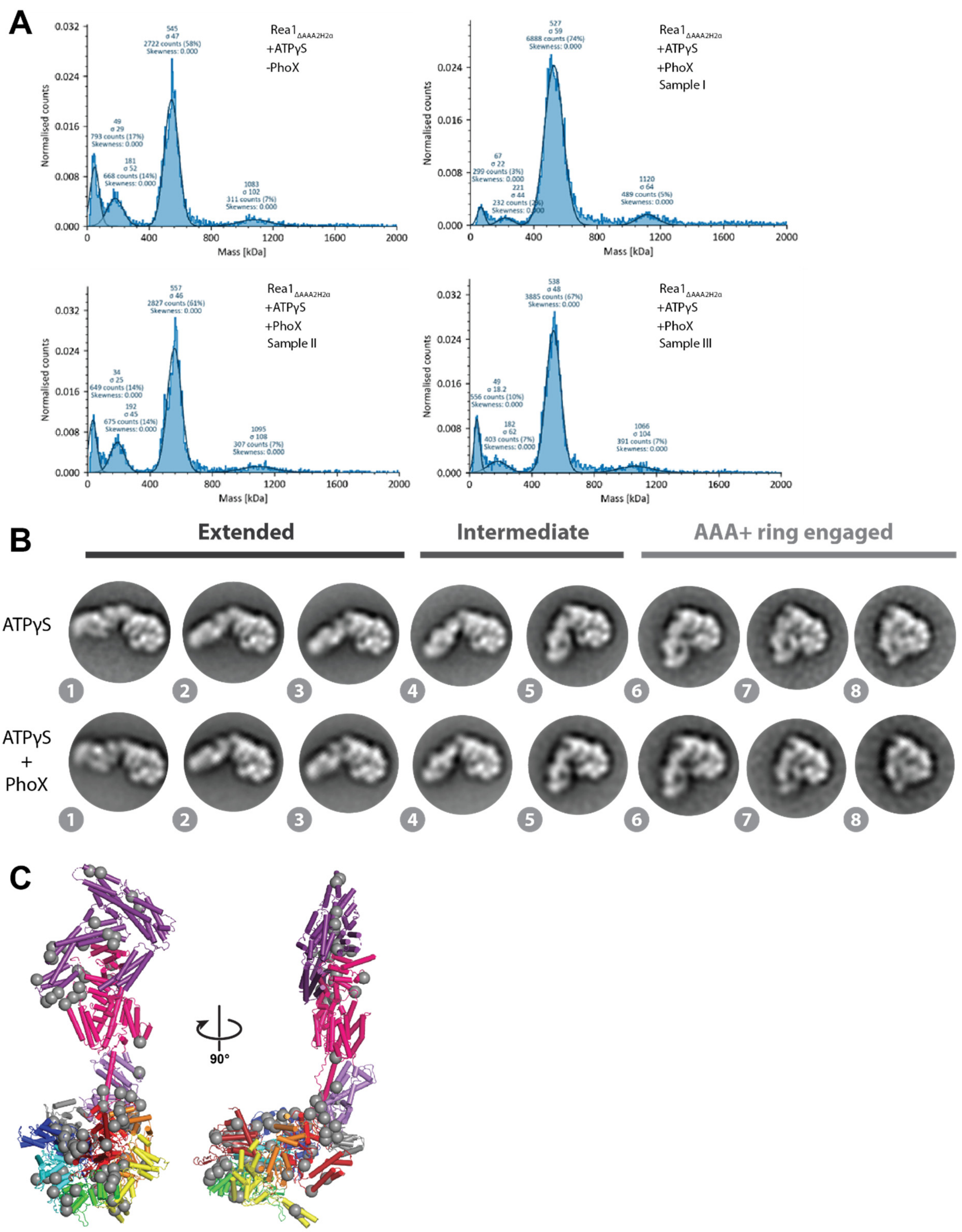


**Supplementary figure 9: Validation of Rea1$_{\Delta AAA2H2\alpha}$ ATPγS PhoX crosslinking. A.** Mass Photometry was used to analyse the amount of unspecifically crosslinked dimers in the sample. The theoretical mass for a Rea1$_{\Delta AAA2H2\alpha}$ monomer is 580 kDa. Solid dark blue lines represent the Gaussian-fits of major species. Compared to the non-crosslinked control (-PhoX) no significant stabilization of Rea1$_{\Delta AAA2H2\alpha}$ dimers is observed in the three analysed PhoX crosslinked samples. **B.** The presence of the PhoX crosslinker does not alter ATPγS induced linker remodelling in Rea1$_{\Delta AAA2H2\alpha}$. **C.** Out of the 183 K-K crosslinks detected in the PhoX crosslinked Rea1$_{\Delta AAA2H2\alpha}$ ATPγS samples, 69 (38%) can be assigned to the straight linker conformation as represented by the Rea1$_{\Delta AAA2H2\alpha}$ ATPγS cryoEM structure (compare Figure 3A), which is expected to be the dominant structural state in the samples. The Cα atoms of crosslinked lysine pairs are highlighted as grey spheres.

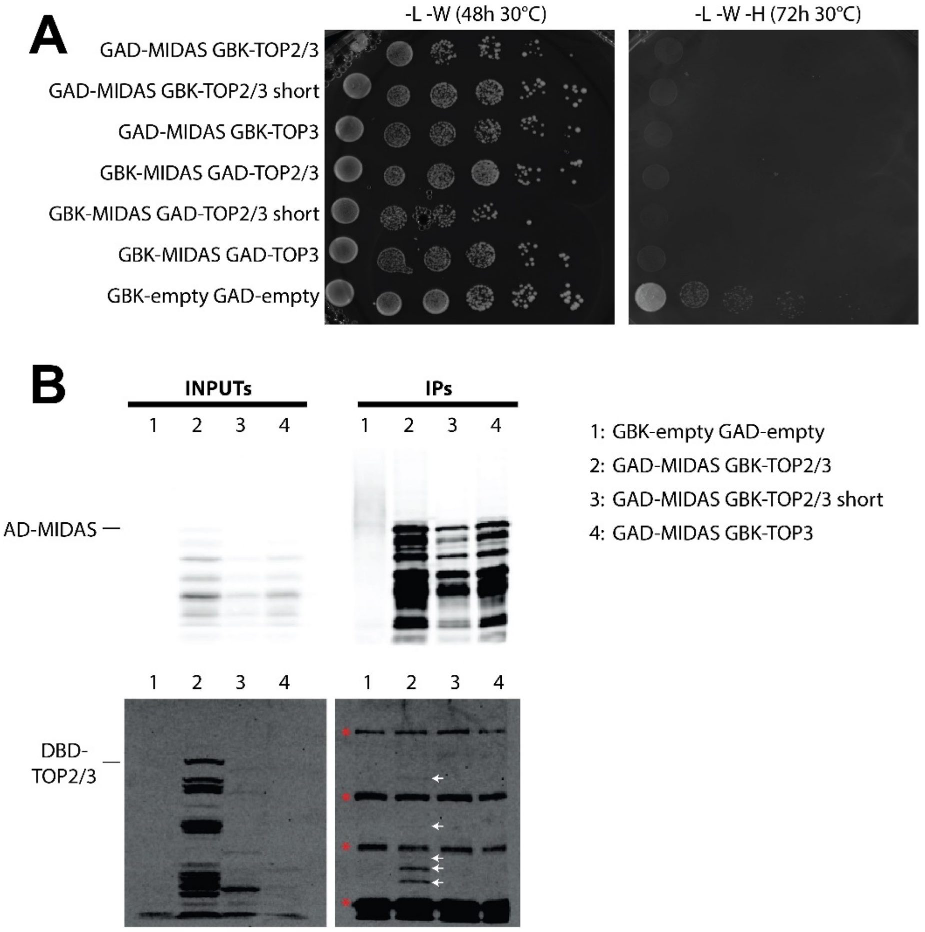


**Supplementary figure 10: The MIDAS domain interacts with the linker top2/top3 region. A.** The MIDAS domain (E4623-S4910) was fused to 3xHA-GAL4AD (GAD, GAL4 activator domain) and the linker top2/top3 region (Y3557-N4041, short: I3601-N4041) or just the top3 region (D3786-E3905) to 3xMyc-GAL4BD (GBK, GAL4 DNA binding domain). Another set of plasmids reversed the GAD/GBK fusion constructs as indicated. The cells grew well on plates selecting the markers of the GAD/GBK plasmids (right panel, plates lacking leucine (L) and tryptophan (W)). Plating the cells on selective medium further lacking histidine (H) to check if the reporter *his3* expression has been induced, revealed that cells expressing the MIDAS/linker top2/top3 constructs grew slower than the empty vector control indicating toxicity to the cells. **B.** In an alternative approach to probe for MIDAS-Linker top2/top3 interactions we carried out GAD-MIDAS immunoprecipitation experiments using anti-HA agarose beads. We checked for expression of the GAD-MIDAS and GBK-Linker top2/top3 constructs in the input lysates by anti-HA and anti-Myc western blot (left upper and left lower panel). The MIDAS and Linker top2/top3 constructs are prone to degradation. The degradation is especially pronounced in the case of the short linker top2/top3 as well as the linker top3 GBK constructs (lanes 3 and 4, left lower panel). Upper right panel: The GAD-MIDAS construct was pulled out from the input lysates via anti-HA agarose beads. The elution was subsequently analysed by anti-HA western blot. GAD-MIDAS can be detected in all elutions (lanes 2 – 4). Lower right panel: The eluates were also analysed by anti-myc western blot to probe for the presence of GBK-Linker top2/top3 constructs. In the case of the GBK-Linker top2/top3 constructs (lane 2), several degradation fragments can be detected (white arrows) indicating that parts of the Linker top2/top3 region are able to interact with the MIDAS domain. Signals marked with red asterisks likely result from detection of anti-HA antibody fragments present in the anti-HA agarose bead elutions by the secondary antibody.

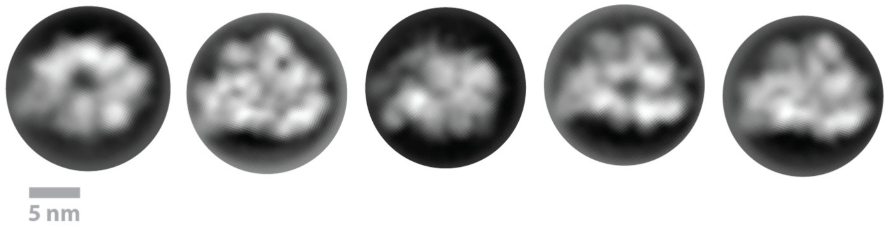


**Supplementary figure 11: Negative stain 2D class averages of Rea1$_{\Delta 3072\text{-}3244}$ in the presence of ATP.** There is no evidence for the linker indicating that the deletion of the globular extension of the middle domain (aa 3072-3244) leads to the degradation of the linker. The degradation of the linker is also supported by SDS-PAGE analysis. The size of the rings in the 2D class averages is comparable with the dimension of the Rea1 AAA+ ring (≈ 12 nm). In the absence of the linker, the AAA+ rings adopt a different orientation on the grid.

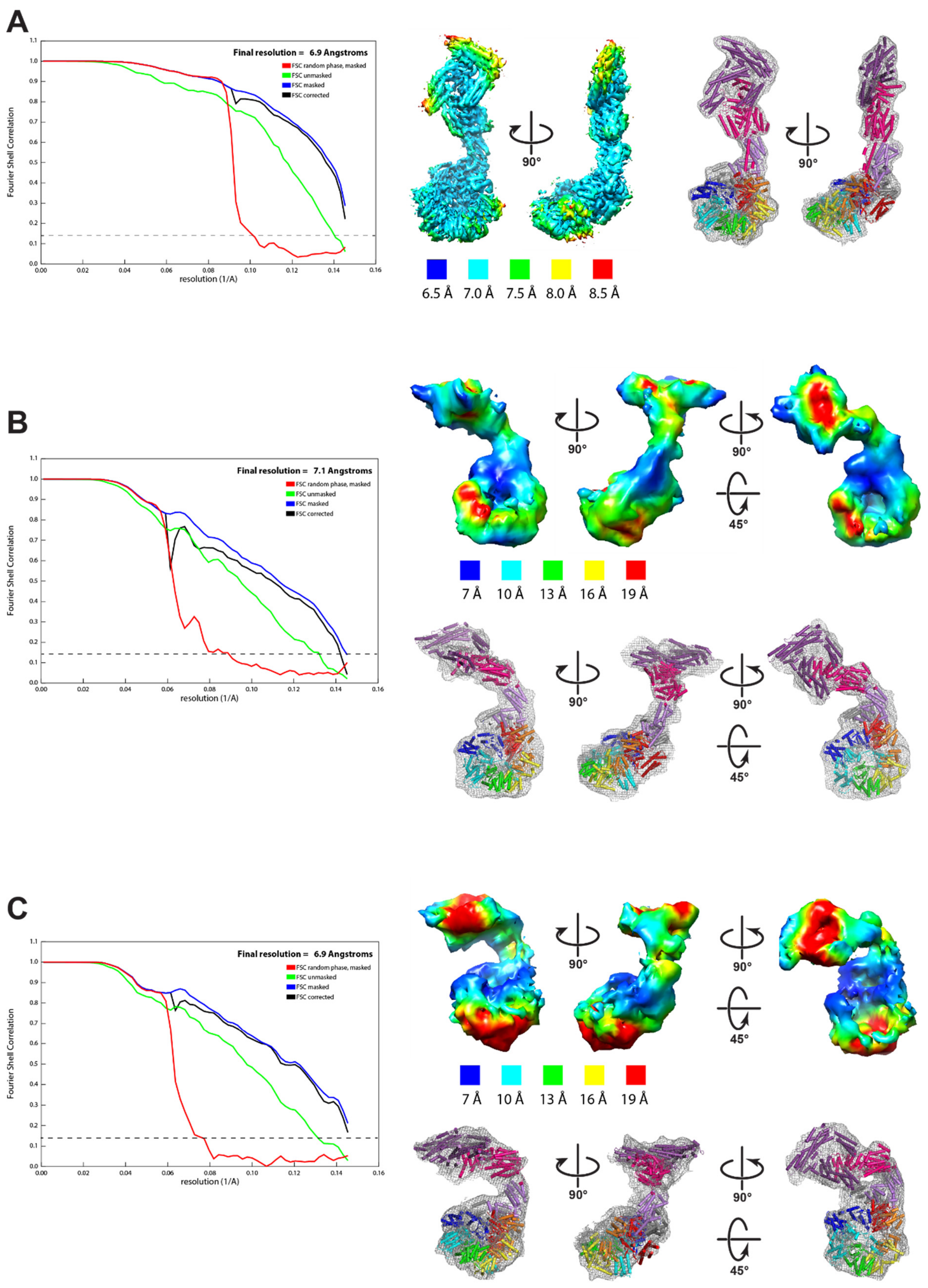


**Supplementary figure 12: Overall quality of the Rea1$_{D2915A-R2976A-D3042A}$ ATP cryoEM maps.** Fourier shell correlation (FSC) plot for half-maps of the 3D reconstructions (left panels, 0.143 FSC criteria is indicated as horizontal dashed line), local resolution maps (middle and right panels) and match of structure in map (right and lower right panels) of **A.** Conformation I, **B.** Conformation II and **C.** Conformation III. In

A. the resolution is of sufficient quality to identify secondary structure elements. In B. and C. the resolution is of sufficient quality to dock in the linker middle-top domains and the linker stem-AAA+ring-NTD. Due to the use of binned data the 0.143 FSC criteria has not been reached in A. and C. No substantial improvements in the resolution are to be expected with the unbinned data.

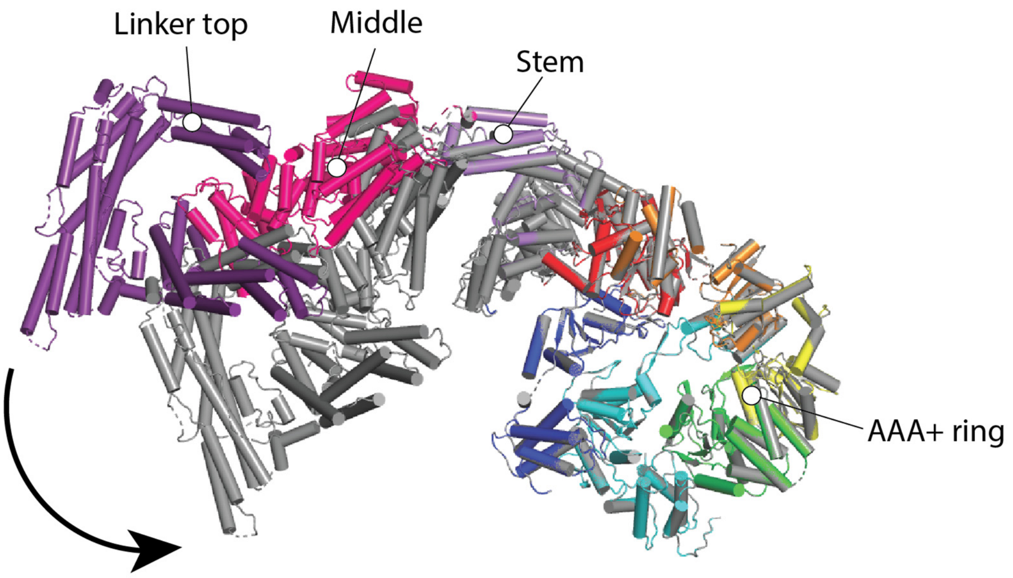


**Supplementary figure 13: Conformations II and III of Rea1$_{D2915A\text{-}R2976A\text{-}D3042A}$ ATP are related by a swing of the linker middle and top domains towards the AAA+ ring.** Conformation II is color coded, Conformation III is shown in grey. The structures have been aligned on the AAA+ rings. The black arrow indicates the swing towards the AAA+ ring. The region between the linker middle and stem domains acts as pivot point.

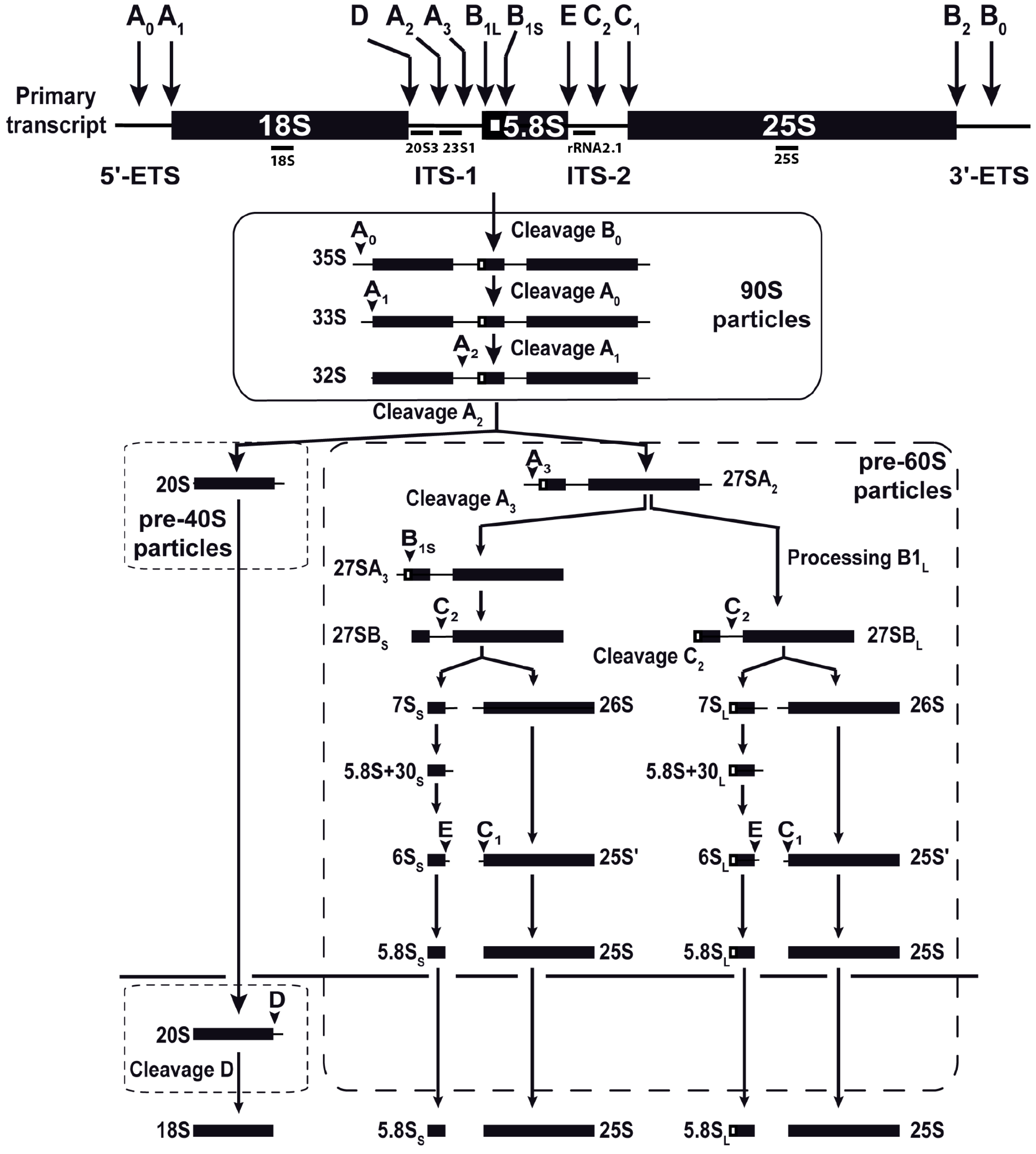


**Supplementary figure 14: Cartoon of the pre-rRNA processing pathway in *S. cerevisiae*.**

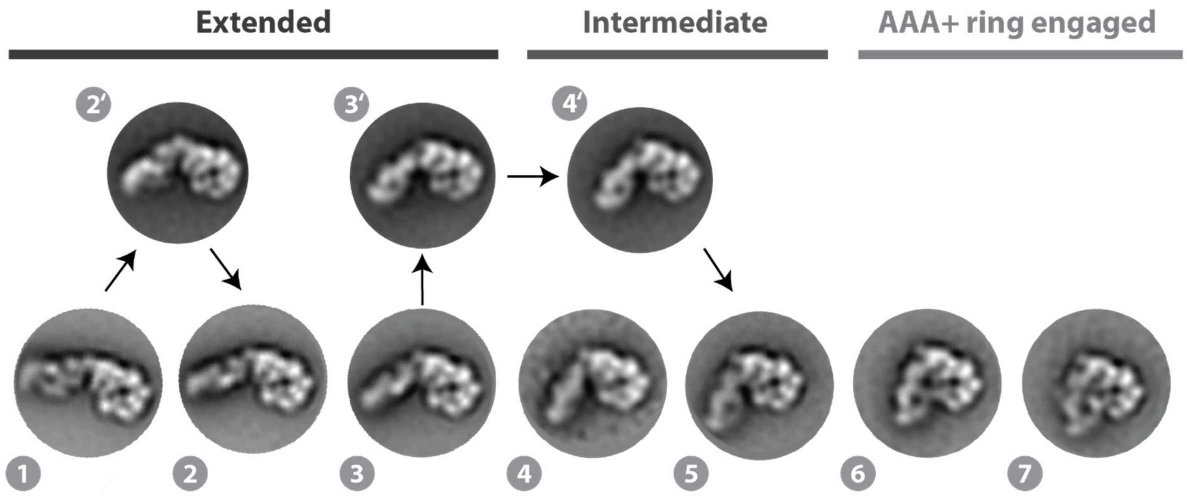


**Supplementary figure 15: The swing and the rotation of the linker top and middle domains during linker remodelling are not strictly correlated.** In addition to the linker states most commonly observed in our data sets (here states 1 – 7 of Rea1$_{wt}$ ATP as an example), additional extended and intermediate linker remodelling states were occasionally detected. Linker state 2’ represents a swing from state 1 towards the AAA+ ring without rotation. In state 3’ the linker middle and top domains are already fully rotated before reaching the proximity of the AAA+ ring. They swing without rotation via state 4’ to state 5.

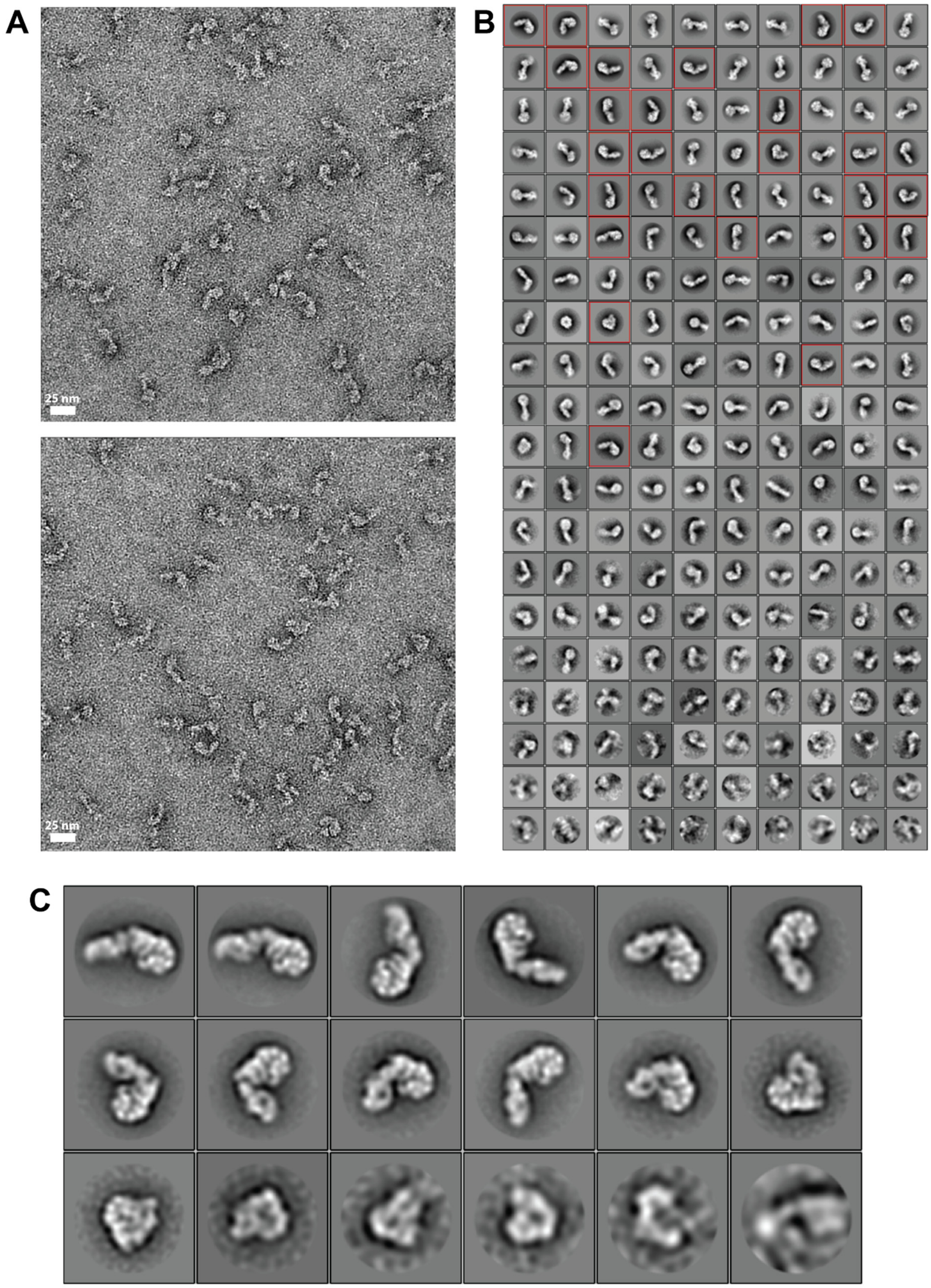


**Supplementary figure 16: Quality of negative stain EM data. A.** Two representative micrographs of the Rea1$_{\Delta AAA2H2\alpha}$ ATPγS data set. **B.** Initial 2D classification on 4x binned particles. 2D classes representing AAA+ ring top views of interest (red squares) were selected for a 2$^{nd}$ round of 2D classification. **C.** 2$^{nd}$ round of 2D classification with the un-binned particles selected in B.
